# Sources of information waste in neuroimaging: mishandling structures, thinking dichotomously, and over-reducing data

**DOI:** 10.1101/2021.05.09.443246

**Authors:** Gang Chen, Paul A. Taylor, Joel Stoddard, Robert W. Cox, Peter A. Bandettini, Luiz Pessoa

## Abstract

Neuroimaging relies on separate statistical inferences at tens of thousands of spatial locations. Such massively univariate analysis typically requires an adjustment for multiple testing in an attempt to maintain the family-wise error rate at a nominal level of 5%. First, we examine three sources of substantial information loss that are associated with the common practice under the massively univariate framework: (a) the hierarchical data structures (spatial units and trials) are not well maintained in the modeling process; (b) the adjustment for multiple testing leads to an artificial step of strict thresholding; (c) information is excessively reduced during both modeling and result reporting. These sources of information loss have far-reaching impacts on result interpretability as well as reproducibility in neuroimaging. Second, to improve inference efficiency, predictive accuracy, and generalizability, we propose a Bayesian multilevel modeling framework that closely characterizes the data hierarchies across spatial units and experimental trials. Rather than analyzing the data in a way that first creates multiplicity and then resorts to a post hoc solution to address them, we suggest directly incorporating the cross-space information into one single model under the Bayesian framework (so there is no multiplicity issue). Third, regardless of the modeling framework one adopts, we make four actionable suggestions to alleviate information waste and to improve reproducibility: 1) abandon strict dichotomization, 2) report full results, 3) quantify effects, and 4) model data hierarchies. We provide examples for all of these points using both demo and real studies, including the recent NARPS investigation.

## 1 Introduction

> *Statisticians classically asked the wrong question* – *and were willing to answer with a lie. They asked “Are the effects of A and B different?” and they were willing to answer “no.”*
>
> *All we know about the world teaches us that the effects of A and B are always different – in some decimal place – for any A and B. Thus asking “are the effects different?” is foolish.*
>
> — John W. Tukey, “The Philosophy of Multiple Comparisons”, Statistical Science (1991)

Functional magnetic resonance imaging (FMRI) is a mainstay technique of human neuroscience, which allows the study of the neural correlates of many functions, including perception, emotion, and cognition. The basic spatial unit of FMRI data is a *voxel* ranging from 1-3 mm on each side. As data are collected across time when a person performs tasks or remains at “rest”, FMRI datasets contain a time series at each voxel. Typically, tens of thousands of voxels are analyzed simultaneously. Such a “divide and conquer” approach through *massively univariate analysis* necessitates some form of multiple testing adjustment via procedures based on Bonferroni’s inequality or false discovery rate.

Conventional neuroimaging inferences follow the null hypothesis significance testing framework, where the decision procedure dichotomizes the available evidence into two categories at the end. Thus, one part of the evidence survives an adjusted threshold at the whole brain level and is considered *statistically significant* (informally interpreted as a “true” effect) while the other part is ignored (often misinterpreted as “not true”) and by convention omitted and hidden from public view.

A recent study^1^ (referred to as NARPS hereafter) offers a salient opportunity for the neuroimaging community to reflect about common practices in statistical modeling and the communication of study findings. The study recruited 70 teams charged with the task of analyzing a particular FMRI dataset and reporting results; the teams simply were asked to follow data analyses routinely employed in their labs at the whole-brain voxel level (but note that nine specific research hypotheses were restricted to only three brain regions). NARPS found large variability in reported decisions, which were deemed to be sensitive to analysis choices ranging from preprocessing steps (e.g., spatial smoothing, head motion correction) to the specific approach used to handle multiple testing. Based on these findings, NARPS outlined potential recommendations for the field of neuroimaging research.

Despite useful lessons revealed by the NARPS investigation, the project also exemplifies the common approach in neuroimaging of generating categorical inferential conclusions as encapsulated by the “significant vs. nonsignificant” maxim. In this context, we address the following questions:

1. Are conventional multiple testing adjustment methods informationally wasteful?
2. The NARPS study suggested that there was “substantial variability” in reported results across teams of investigators studying the same dataset. Is this conclusion dependent, at least in part, on the practice of drawing inferences binarily (i.e., “significant” vs. “non significant”)?
3. What changes can the neuroimaging field make in modeling and result reporting to improve replicability?

In this context, we consider inferential procedures not strictly couched in the standard null hypothesis significance testing framework. Rather, we suggest that multilevel models, particularly when constructed within a Bayesian framework, provide powerful tools for the analysis of neuroimaging studies given the data’s inherent hierarchical structure. As our paper focuses on hierarchical modeling and dichotomous thinking in neuroimaging, we do not discuss the broader literature on Bayesian methods applied to FMRI^2^.

## 2 Massively univariate analysis and multiple testing

We start with a brief refresher of the conventional statistical framework typically adopted in neuroimaging. Statistical testing begins by accepting the null hypothesis but then rejecting it in favor of the alternative hypothesis if the current data for the effect of interest (e.g., task A vs. task B) or potentially more extreme observations are unlikely to occur under the effect is zero. Because the basic data unit is the voxel, one faces the problem of performing tens of thousands of inferences across space *simultaneously.* As the spatial units are not independent of one another, adopting an adjustment such as Bonferroni’s is unreasonably conservative. Instead, the field has gradually settled into employing a cluster-based approach: what is the size of a spatial cluster that would be unlikely to be observed under the null scenario?

Accordingly, a two-step procedure is utilized: first threshold the voxelwise statistical evidence at a particular (or a range of) voxelwise *p*-value (e.g., 0.001) and then consider only contiguous clusters of evidence (Fig. 1). Several adjustment methods have been developed to address multiple testing by leveraging the spatial relatedness among neighboring voxels. The stringency of the procedures has been extensively debated over the past decades, with the overall probability of having clusters of a minimum spatial extent given a null effect estimated by two common approaches: a parametric method^3,4^ and a permutation-based approach^5^. For the former, recent recommendations have resulted in the convention of adopting a primary threshold of voxelwise *p* = 0.001 followed by cluster size determination^6,7^; for the latter, the threshold is based on the integration between a range of statistical evidence and the associated spatial extent^5^.

**Figure 1:**
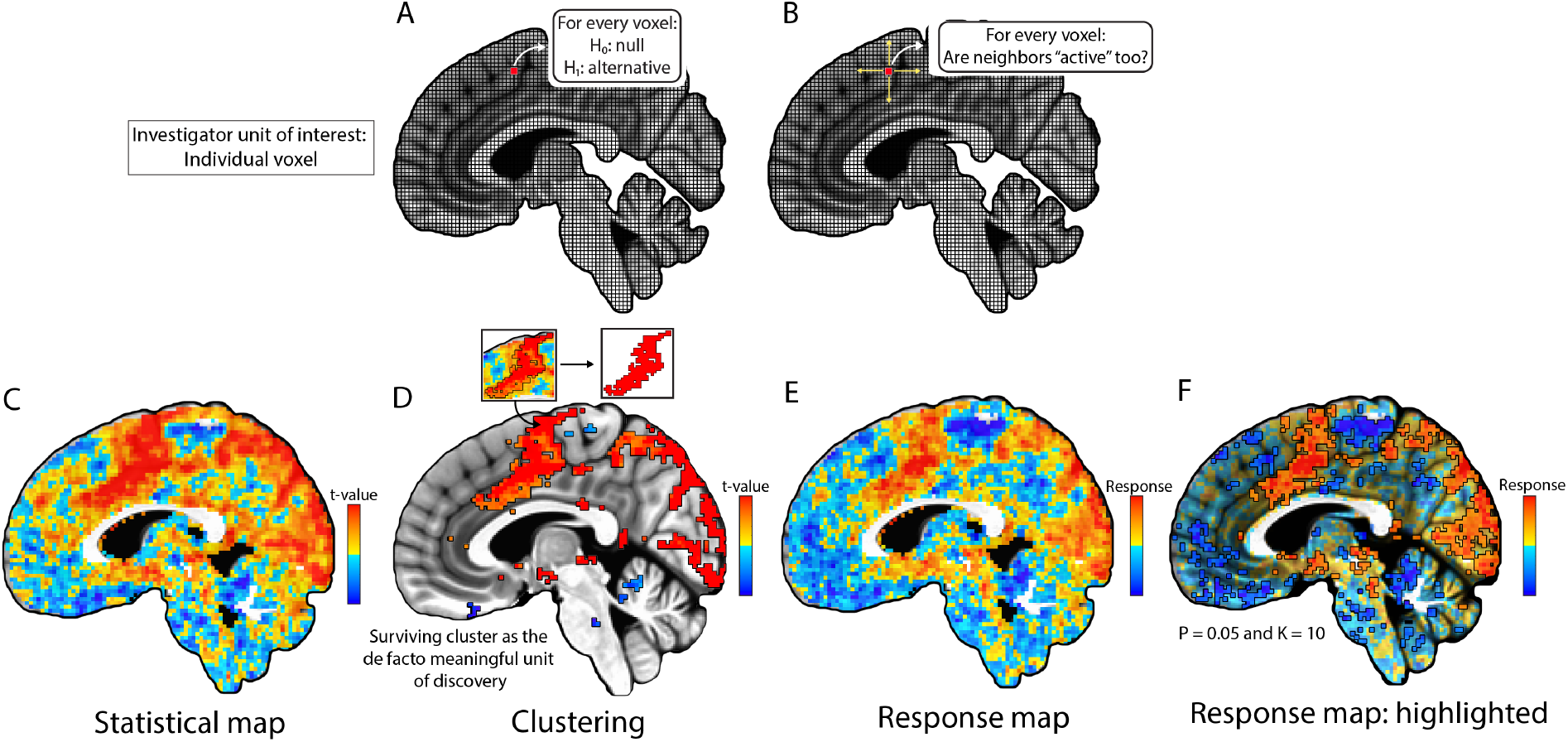
Statistical inferences in neuroimaging. (A) Schematic view of standard analysis: each voxel among tens of thousands of voxels is tested against the null hypothesis (voxel not drawn to scale). (B) Clusters of contiguous voxels with strong statistical evidence are adopted to address the multiple testing problem. (C) Full statistical evidence for an example dataset is shown without thresholding. (D) The statistical evidence in (C) is thresholded at voxelwise *p* = 0.001 and a cluster threshold of 20 voxels. The left inset shows the voxelwise statistical values from (C) while the right inset illustrates the surviving cluster. (E) The map of effect estimates that complements the statistical values in (C), providing percent signal change or other index of response strength, is shown. (F) For presenting results, we recommend showing the map of effect estimates, while using the statistical information for little or moderate thresholding: “highlight” parts with strong statistical evidence, but do not “hide” the rest.

### 2.1 Problems of multiple testing adjustments

At least four limitations are associated with multiple testing adjustments leveraged through spatial extent^8^.

1. *Conceptual inconsistency*. Consider that the staples of neuroimaging research are the maps of statistical evidence and associated tables. Both typically present only the statistic (e.g., *t*) values. However, this change of focus is inconsistent with cluster-based inference: after multiple testing adjustment the proper unit of inference is the cluster, not the voxel. Once “significant” clusters are determined, one *should* only speak of clusters and the voxels inside each cluster *should* no longer be considered meaningful inferentially. In other words, the statistical evidence for each surviving cluster is deemed at the “significance” level of 0.05 and the voxelwise statistic values lose direct interpretability. Therefore, voxel-level statistic values in brain maps and tables in the literature should not be taken at face value. Furthermore, as a cluster is purely defined through statistical evidence, it is usually not aligned with any anatomical region, presenting a spatial specificity problem. To resolve the issue, the investigator typically reduces the cluster to a “peak” voxel with the highest statistical value and uses its location as the evidence for the underlying region. A conceptual inconsistency results from these two transitional steps: one from a cluster to its peak voxel, and then another from the voxel to an anatomical region. Furthermore, when a cluster spans over more than one anatomical region, no definite solutions are available to resolve the inferential difficulty. Although this issue of conceptual inconsistency and spatial specificity has been discussed in the past^7,8^, it remains underappreciated, and researchers commonly do not adjust their presentations to match the cluster-level effective resolution.
2. *Heavy penalty against small regions*. With the statistical threshold at the spatial unit level traded off with cluster extent, larger regions might be able to survive with relatively weaker statistical strength while smaller regions would have to require much stronger statistical strength. Therefore, multiple testing adjustments always penalize small clusters. Regardless of the specific adjustment method, anatomically small regions (e.g., those in the subcortex) are intrinsically disadvantaged even if they have the same amount of statistical evidence. In other words, ideally the evidence for a brain region should be assessed solely in light of its effect strength, not dependent of its anatomical size; thus, the conventional multiple testing adjustment approaches are unfair to small regions because of its heavy reliance on spatial relatedness among contiguous neighborhood.
3. *Sensitivity to data domain*. As the penalty for multiplicity becomes heavier when more spatial units are involved, one could explore various surviving clusters by changing the data space (e.g., “small volume correction”), resulting in some extent of arbitrariness: one cluster may survive or fail depending on the spatial extent of the data. Because of this vulnerability, it is not easy to draw a clear line between a justifiable reduction of data and an exploratory search (e.g., “small volume correction”).
4. *Difficulty of assigning uncertainty.* As the final results are inferred at the cluster level, there is no clear uncertainty that can be attached to the effect magnitude at the cluster level. On the one hand, a cluster either survives or not under a dichotomous decision. On the other hand, due to the interpretation difficulty of voxel-level statistical evidence, it remains challenging to have, for example, a standard error (“error bar”) associated with the average effect at the cluster level.

It is worth remembering a key goal of data processing and statistical modeling: to take a massive amount of data that is not interpretable in its raw state, and to extract and distill meaningful information. The preprocessing parts aim to reduce distortion effects, where as statistical models intend to account for various effects. Overall, there is a broad trade-off along the “analysis pipeline”: we increase the digestibility of the information at the cost of reducing information. Fig. 2A illustrates these key aspects of the process of information extraction in standard FMRI analysis. The input data of time series across the brain for multiple participants are rich in information, but of course not easily interpretable or “digestible.” After multiple preprocessing steps followed by massively univariate analysis, the original data are condensed into two pieces of information at each spatial unit: the point estimate and the standard error. Whereas this process entails considerable reduction of information, it produces usefully digestible results; we highlight this trade-off in Fig. 2B. Here, “information” refers broadly to the amount and content of data present in a stage (e.g., for the raw data, the number of groups, participants, time series lengths, etc.). “Digestibility” refers to the ease with which the data are presentable and understandable (e.g., two 3D volumes vs one; a 3D volume vs a table of values). Following common practice, many investigators discard effect magnitude information to focus on summary statistics, which are then used to make binarized inferences by taking into account multiple testing. These steps certainly aid in reporting results and summarizing potentially some notable aspects of the data. However, below, we argue that the overall procedure leads to information waste, and that the gained digestibility is relatively small (in addition to generating problems when results are compared across studies). Whereas we focus our discussion on whole-brain voxel-based analyses, similar issues apply in other types of analysis for region-based and matrix-based data.

**Figure 2:**
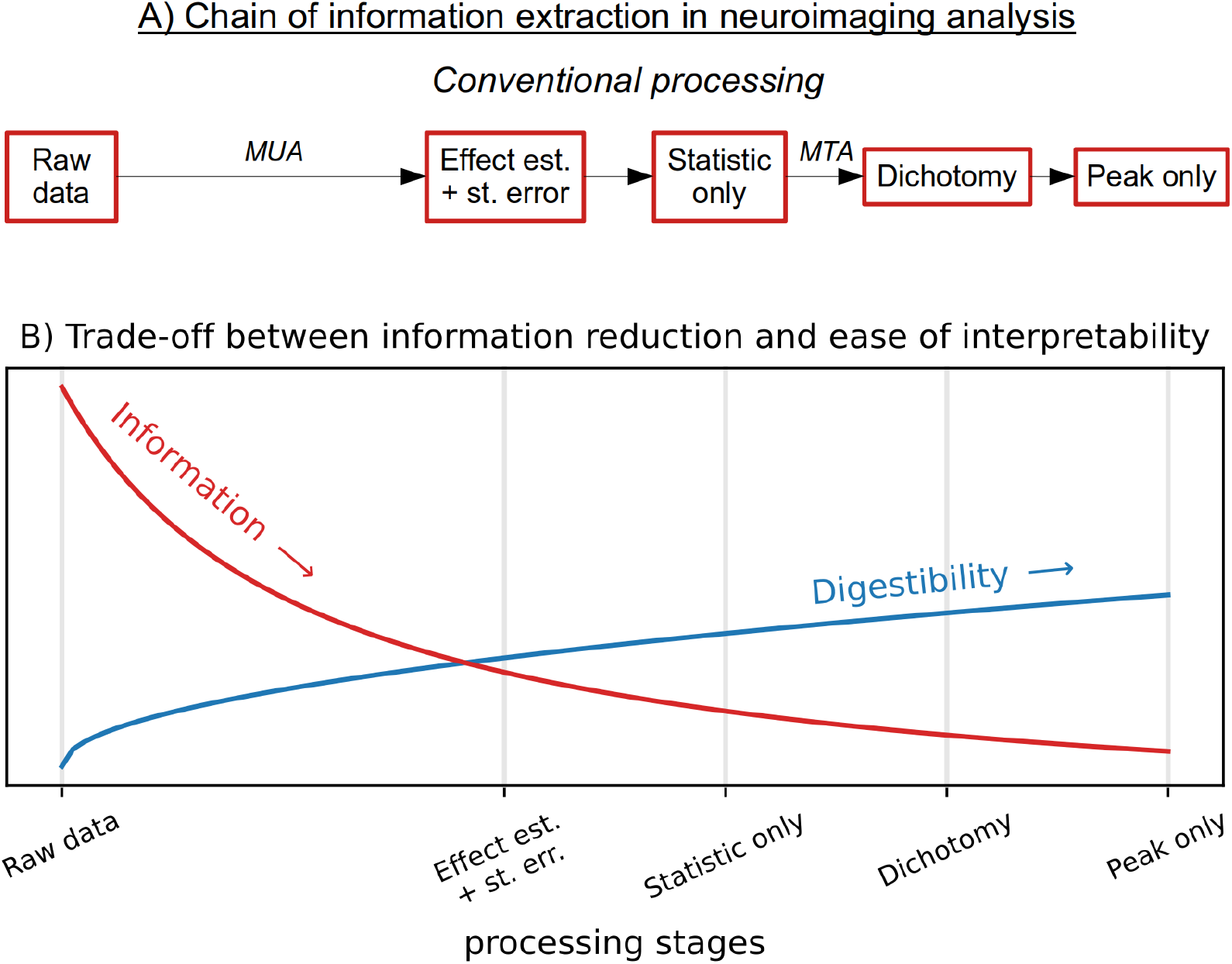
A schematic of conventional information extraction in neuroimaging. (A) The processing chain starts with raw data. Massively univariate analysis (MUA) produces a point estimate and its uncertainty (standard error) at every spatial unit. These are reduced to a single statistic map, which is then dichotomized using thresholding through multiple testing adjustment (MTA); finally, the analyst summarizes the regions based solely on their peak values, ignoring spatial extent. (B) The inherent trade-off between “information” and “digestibility” (*y*-axis has arbitrary units). While summarizing peak locations of dichotomized regions is a highly digestible form of output, this also entails a severe information loss. Here, we argue that providing effect estimates and standard errors, if possible, would be preferable, striking a better balance between information loss and interpretability.

### 2.2 The implicit assumption of massively univariate analysis

Massively univariate analysis, by definition, models all voxels simultaneously with the assumption that all voxels (typically covering the entire brain) are unrelated to one another and that they do not share information. As a corollary, this also assumes that all possible effects have the same probability of being observed, which is to say that the effects follow a uniform distribution from −∞ to +∞ (Fig. 3A), at times discussed as the *principle of indifference*^9^ or the *principle of insufficient reason*^10^. Adopting this “indifference” approach might be reasonable, especially when the distribution of effects is unknown. However, it may result in information loss and lead to costly statistical accommodations.

**Figure 3:**
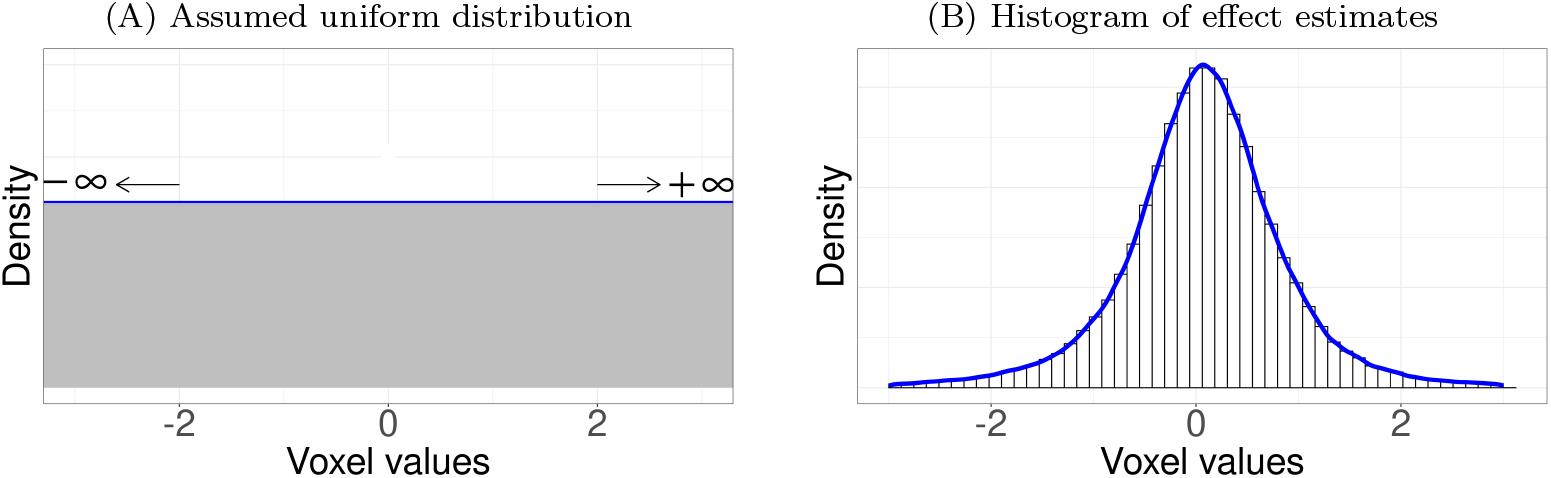
Distributions of effects (“activation strength” in percent signal change) across space. (A) In massively univariate analysis, effects across all spatial units (voxels) are implicitly assumed to be drawn from a uniform distribution. Accordingly, the effect at each spatial unit can assume any value within (−∞, +∞) with equal likelihood. (B) Histogram of effect estimates (percent signal change) across 153,768 voxels in the brain from a particular study. Contrary to the assumption of uniform distribution implicitly made in massively univariate models, the effects approximately trace a Gaussian (or Student’s *t*) distribution.

In this context, we ask the following question: Do FMRI effects across the brain actually follow a uniform distribution, as tacitly assumed in massively univariate analysis, or are they closer to a symmetric bell-shaped distribution? We suggest that a better starting point would be a Gaussian (or possibly something with heavier tails, like Student’s *t*) distribution (Fig. 3B). Conceptually, a Gaussian distribution is a reasonable choice if the effects track an average while also exhibiting a certain extent of variability.

Two direct consequences of massively univariate analysis are information waste and overfitting. Under the principle of insufficient reason, one trusts the *local* “unbiased” point estimates while adjusting the extent of statistical evidence among neighboring spatial units during multiple testing adjustment; however, the loss of modeling efficiency and accuracy at the global level can only be partly recouped at the neighborhood, not *global,* level^8^. In addition to potentially excessive penalties due to information waste, the principle of indifference has another important ramification: *overfitting.* As spatial units are treated as parallel entities – not part of the data hierarchy – in the model, global information shared across space is not leveraged and calibrated, leading to the loss of modeling efficiency. In other words, under massively univariate analysis, the model is free to fit the voxel’s data in any way it can as all possible effect magnitudes are equally likely. As the field of machine learning has demonstrated repeatedly, overfitting is a serious problem because of compromised generalizability (is it possible to learn from a sample to predict out-of-sample test cases?). Thus, whereas the massively univariate approach offers unbiased estimates at the spatial unit level (via least squares or maximum likelihood), it tends to fit individual voxels overly close to the sample data at hand. Consequently, it may lead to a suboptimal tradeoff between bias and variance and pay the cost of overfitting the data with reduced predictive accuracy when future data are considered.

What can be done to address the issues of information waste and overfitting? As a first step, we suggest that voxelwise modeling should take a holistic view, considering the effects as distributed normally (or according to Student’s *t*). The reasoning here is analogous to when we assume that effects are normally distributed *across subjects* (or “random-effects” in linear mixed-effects modeling) in neuroimaging studies, allowing inferences at the population level. In a similar fashion, we propose conceptualizing voxel-level effects in terms of sampling from a normally distributed hypothetical pool of effects, instead of adopting the stance of complete ignorance (i.e., uniform distribution).

Technically, we can say that the effect distribution across spatial units, 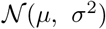, forms a *prior distribution* in the Bayesian sense where the two hyperparameters, the mean *μ* and the standard deviation *σ*, are basically estimated from the data. On the one hand, the variability of the data across spatial units (see Fig. 3B) determines the magnitude of *σ*. On the other hand, the estimated *σ* influences the estimates of *μ* across the spatial units through a process of “information sharing”, regularization or *partial pooling*. For example, if most of the individual effects across space are estimated to be small and close to zero, *σ* is estimated to be small, which further tends to decrease the individual effects, a situation also referred to as *shrinkage*.

We do not claim that the conventional approach is not valid. Instead, we suggest that the indifference assumption is an inefficient way of modeling the data, which can benefit from information sharing across space. Note that when NARPS summarized team results to make meta-analytic statements, they did not assume a uniform distribution of effects across teams; instead, they assumed that the results across studies would follow a Gaussian distribution. In other words, they did not treat the teams as “isolated trees”. Interestingly, they did not adjust for multiple testing when interpreting individual team inferences, even though 70 teams simultaneously analyzed the data and provided separate results. We agree that the adoption of a Gaussian prior is a sensible approach: it assumes that the results track an average population effect, while exhibiting variability across teams. However, we propose that such utilization of priors does not have to be limited to or stopped at meta-analysis across different analytical pipelines; rather, information integration through a “forest perspective” can be equally applied to modeling across all hierarchies, including the levels of voxels, regions, experimental trials, and participants.

## 3 Problems of dichotomous thinking

Data compression is essential in science so that complex information originating from large datasets can be encapsulated in terms of key findings (Fig. 2). Nevertheless, we believe that neuroimaging’s common practice of adhering to multiple testing adjustments together with dichotomization (“significant or not”) is detrimental to scientific progress. Take the process of examining the results by first insisting on the use of a cluster-based approach through a strict voxelwise threshold (*p* < 0.001) coupled with a minimum cluster extent (say, 50 voxels). In many instances, the analyst will miss the opportunity to make important novel observations; maybe some non-surviving clusters are just over 30 voxels (not to mention 49 voxels), for instance. The permutation-based approach to handling multiplicity suffers from the same issue.

In the last decade, statisticians and practitioners have extensively discussed pervasive issues with the practice of significance testing^11^. As typically practiced in neuroimaging, solely focusing on and reporting statistical results that have survived significance filtering leads to issues such as overestimation (“winner’s curse”, publication bias^12,13^ or type M error^14^) and type S error (incorrect sign)^14^, A widespread problem is the disconnect between null hypothesis significance testing and the way investigators think of their research hypothesis. The *p*-value is the probability (or the extent of inconsistency or “surprise”) of a random process generating the current data or *potentially more extreme observations* if a null effect were actually true (conditioned on the experimental design, the adopted model, and underlying assumptions). In contrast, an investigator is likely more interested in the probability of a research hypothesis (e.g., a positive effect) given the data. Misinterpretations of the *p*-value frequently lead to conceptual confusions^15^. The *p*-values are also affected by the extent to which the model in question and its assumptions are suited for the data at hand.

Recognizing deep and entrenched research practices, the American Statistical Association has issued guidelines and proposed potential reforms^16^. In our view, this important debate has not penetrated the neuroimaging community sufficiently. Given the expense and risk of collecting FMRI data, it is important to embrace methods that address problems with “significance testing” while simultaneously decreasing information waste. In a nutshell, we believe experimental science and discovery is a highly complex process that cannot be simplified and reduced to drawing a sharp line with the use of thresholding procedures, regardless of their numerical stringency and formal mathematical properties.

Problems with boiling down complicated study designs into binary decisions are further aggravated by the empirical observation that, as discussed, effects across the brain tend to follow a Gaussian distribution (Fig. 3B). Consistent with this notion, one study reported that over 95% of the brain was engaged in a simple visual stimulation plus attention task when large participant samples were considered^17^. In contrast, most studies in neuroimaging only report a few brain regions that happen to survive the artificial dichotomization based on the currently accepted spatial adjustment criteria. The large gap between the engagement in most regions and the few regions reported in the literature is likely due to limited sample sizes as well as to information waste. More generally, many domains of research appear to be characterized by a very large number of “small effects” as opposed to a few “large effects”, including genetics^18,19^ and most likely brain research itself. Thus, a data analysis framework, such as null hypothesis significance testing, that seeks to binarize results only using statistical evidence (while ignoring separate effect estimates and uncertainties) is potentially problematic. We conjecture that this could represent the case in neuroimaging, where effects are present across large numbers of spatial units (voxels or brain regions) at varying strengths. In addition, it is worth noting that, even though family-wise error rate is the major leverage adopted to control multiplicity in neuroimaging, the arbitrariness issue involved in dichotomization equally applies to other notions such as false discovery rate.

We propose that a more productive approach is to refocus research objectives away from trying to uncover “real” effects. Specifically, more emphasis can be placed on discussing effects with stronger evidence, comparing large against small ones, or effects with smaller uncertainty against ones with larger uncertainty (Fig. 4, right). Accordingly, methodological research goals should concentrate on developing an efficient experimental design and improving statistical modeling. More broadly, we advocate for approaches that are more accepting of the statistical uncertainty associated with data analysis, that is, more cognizant of inherent variability in data. In particular, investigators should not treat results that survive a particular threshold as “real” with the rest as “non-effects”, and thus should not describe effects that survive as “facts”. In other words, we recommend that one should avoid a result description with definitive certainty (e.g., null effect); even the typical language of “active/activated voxels/regions” comes with substantial perils. In general, we encourage further discussion about better and more nuanced ways of summarizing research findings.

**Figure 4:**
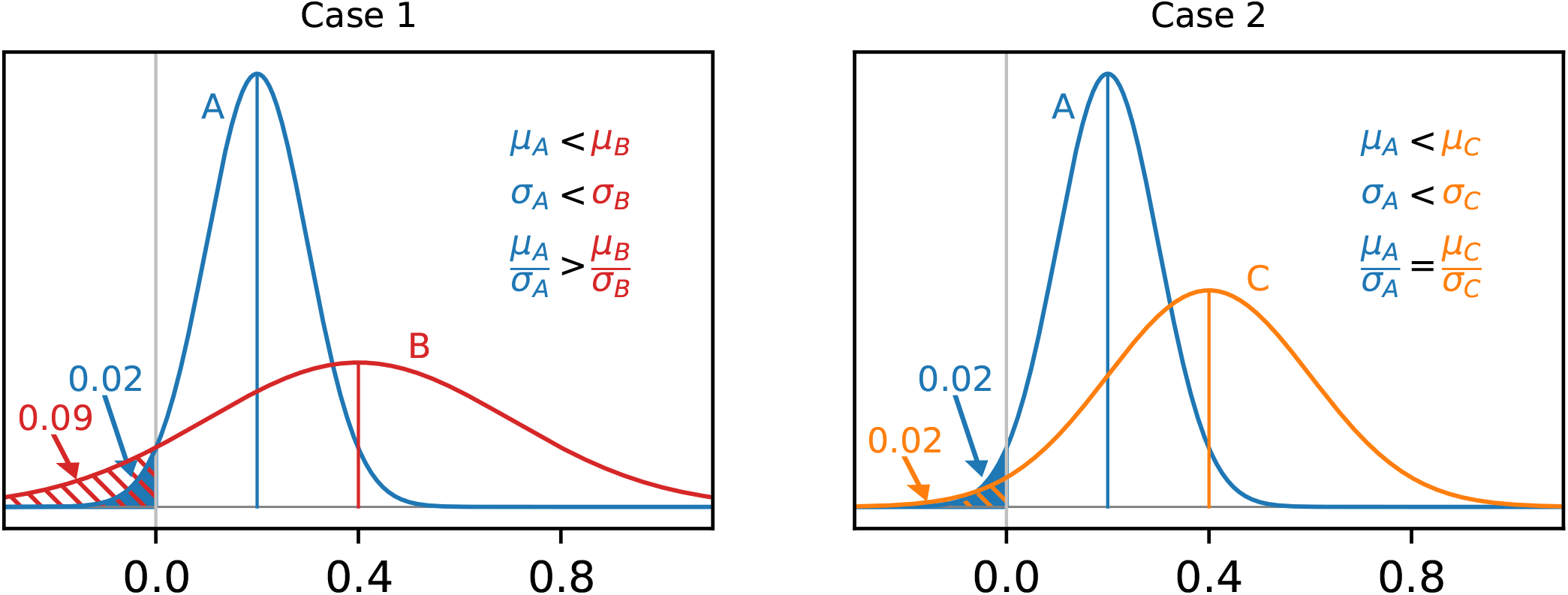
Implications of dichotomization in conventional statistical practice. (Left) What is the difference between a statistically significant result and one that does not cross threshold? Between the two hypothetical effects *A* and *B* that independently follow 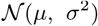 (*μ_A_* = 0.2, *σ_A_* = 0.1 (blue); *μ_B_* = 0.4, *σ_B_* = 0.3 (red)), only A would be considered statistically significant. As the difference between the two random variables associated with *A* and *B* follows 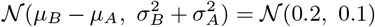, the difference is not considered statistically significant (*p* = 0.26, area under the density of 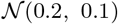 on the left side of *y*-axis), and effect *B* is mostly larger than *A* with a probability of 0.74 (area under the density of 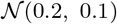 on the right side of *y*-axis). (Right) How much information is lost due to the focus on binary statistical decisions? The two hypothetical effects *A* and *C* (independently with 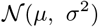: *μ_A_* = 0.2, *σ_A_* = 0.1 (blue); *μ_C_* = 0.4, *σ_C_* = 0.2 (orange)) have the same *t*- and *p*-values, and would be deemed indistinguishable in terms of statistical evidence alone. However, as the difference between the two random variables associated with *A* and *C* follows 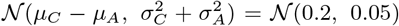, *C* is mostly larger than *A* with a probability of 0.81 (area under the density of 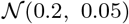 on the right side of *y*-axis). This comparison illustrates the information loss when the sole focus is on statistic or *p* value, which is further illustrated between the second and third blocks in Fig. 2.

### 3.1 Neglect of effect magnitude and uncertainty measures

Statistical significance combines two underlying pieces of information: the effect estimate and its uncertainty. However, because statistical significance is used as a filtering mechanism, investigators typically do not emphasize the “uncertainty” component, even though the underlying machinery is of probabilistic nature. As a result, in practice a statistically significant result tends to be treated as “real, with zero uncertainty”. In addition, a nonsignificant result is often interpreted as showing the absence of an effect, as opposed to representing the lack of sufficient evidence to overturn the null hypothesis, despite repeated warnings against such conclusions in statistical textbooks and training. While these two issues are interpretational problems, they occur so often with the null hypothesis significance testing paradigm that they have almost become part of the paradigm itself, making it easy to fall into these conceptual traps.

Some of the above issues can be illustrated by considering the NARPS study. Given the findings from the 70 independent teams, NARPS performed two types of meta-analysis: one with binarized team reports (logistic regression), and another solely based on statistical values. In the binarized case, the result of each individual team was considered either present (value of 1: the presence of strong evidence is interpreted as an evidence with no uncertainty) or absent (value of 0: the absence of evidence is equated to an evidence of absence). NARPS simply interpreted their meta-analytic findings as indicating substantial variability in study results across different analytical pipelines. A well-known problem with the dichotomization approach is that it treats *p*-values of 0.049 and 0.051, for example, as categorically distinct. On the one hand, the difference between a statistically significant result may not significantly different from a statistically insignificant one (Fig. 4, left). On the other hand, possibly less appreciated is the fact that the approach neglects differences between the two results that are deemed significant (i.e., in both cases *p* < 0.05) (Fig. 4, right): two results with the same amount of statistical evidence may have nontrivial differences in both effect magnitude and uncertainty. These examples illustrate the extent of information loss due to the sole emphasis on statistical evidence while deemphasizing effect magnitude as routinely practiced in neuroimaging.

To further appreciate the above issues, consider the hypothetical scenario illustrated in Fig. 5. The example could refer to a series of studies that investigated a specific experimental paradigm in the past (e.g., activation in the amygdala due to fearful and neutral faces), or to the case considered by NARPS in which different teams analyzed the same dataset. In the scenario, 3 out of 11 results survive the conventional threshold cutoff (Fig. 5A); one may claim poor reproducibility and “sizeable variation” across individual results, and question the statistical evidence provided by the suprathreshold studies. This situation only worsens if one imposes an adjustment for multiple testing to the statistical threshold due to having 11 parallel inferences: with an adjustment applied, none of the studies would survive dichotomization.

**Figure 5:**
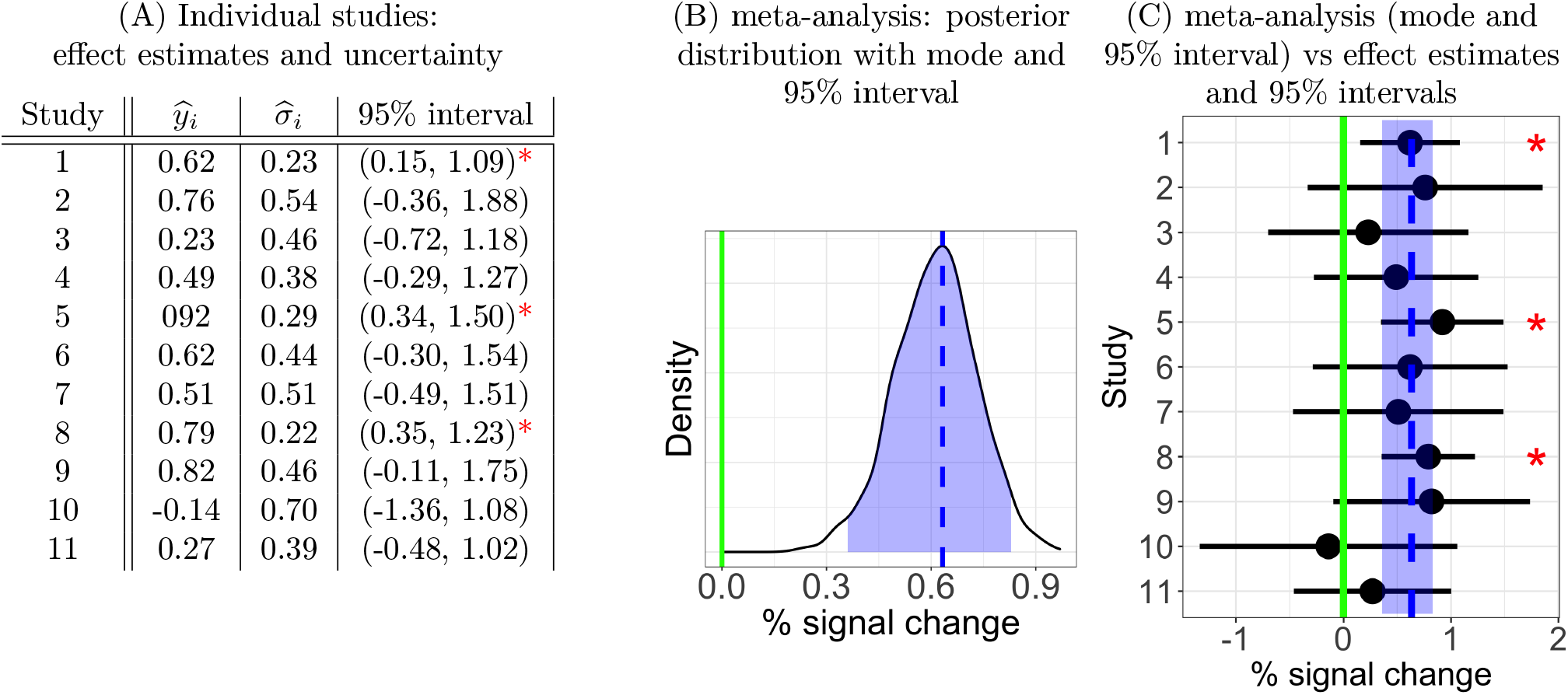
meta-analysis example. (A) Hypothetical results of 11 studies analyzing the same data (or 11 studies of the same task), with results summarized by the estimate of the effect, 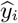 (where *i* is the study index), and its standard error, 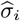. A total of 3/11 effects would be deemed statistically significant (red asterisk) according to standard cutoffs. From this perspective, one might say there is inconsistency or “considerable variability” of study results. (B) A different picture emerges if the same studies are combined in a meta-analysis: the overall evidence (area under the curve the right of zero) points to a positive effect. The posterior distribution of the effect based on Bayesian multilevel modeling provides a richer summary of the results than (A). The shaded blue area indicates the 95% highest density interval (0.36, 0.83) surrounding the mode 0.63 (dashed blue line). (C) The individual results from (A) are presented (dots indicate 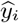, horizontal lines show 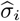, and red asterisks indicate the individually significant studies), along with the meta-analysis distribution information (colors as in B). With the full information present, we can evaluate the study consistency and overall effect more meaningfully.

Instead of a logistic regression based on binarized assessments, an integrative meta-analysis can be performed by combining the full results of *both* the effect estimate and uncertainty from each study. Let us assume that the effect estimates, 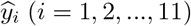, are normally distributed 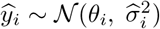 with mean *θ_i_* and variance 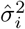. In addition, suppose that the effects themselves, *θ_i_*, follow 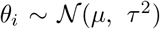 with mean *μ* and variance *τ*^2^. The latter distribution specifies a prior and provides some information to the process, but only minimally: it assumes that the effects *θ_i_* tend to have a bell-shaped, not uniform, distribution, with some values more likely than others. Under this modeling perspective^a^, we obtain a posterior distribution of *μ* (Fig. 5B) with an average effect 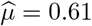 and a 95% uncertainty interval (0.34, 0.85). When this posterior uncertainty interval is reviewed together with the estimates and uncertainties of the 11 individual studies (Fig. 5C), we now have a convenient way to check and evaluate the consistency of the studies; the fact that majority of the individual effect mean values fall within (or just outside) the meta-analysis’s 95% interval indicates a large degree of consistency, rather than a dichotomized assessment with 3 out of 11 “statistically significant” results.

The last result leads to a very different conclusion than when the meta-analysis was based only on binarized statistics, because the proposed analysis uses both the effect estimate and uncertainty of each individual result. Note that the binarized version is highly sensitive to the definition of “significance” used for the individual studies, as well as to the specific adjustment for multiple testing. Clearly, there is considerable information loss in the processes of binarization and multiple testing adjustment. As an alternative, consider having access only to a summary statistic (e.g., Student’s *t*) for each study. A statistic is in essence the ratio of the estimated effect relative to its variability and reduces the two independent pieces of information into one. Whereas including statistic values is a step in the right direction, it is an insufficient one. Displaying both the effect estimate and its variability would provide richer information than a statistic value alone. To see this, consider the simple meta-analysis model described above, where the overall effect estimate for *n* studies, given *τ*, can be stated as^20,21^

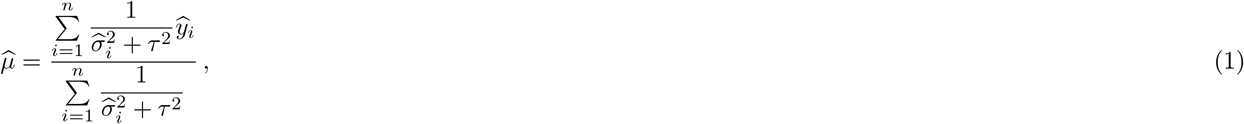

with a standard error 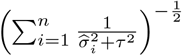. In other words, the full results of the n studies are combined through the weighted average among the individual effects 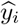 with their associated variances 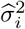, together with the cross-study variance *τ*^2^, inversely playing the role of weighting.

The preceding meta-analysis illustrates the value of reporting *both* effect estimate *and* uncertainty values in scientific communication. As FMRI signals do not follow a ratio scale with a true zero, we recommend reporting percent signal change or another index of magnitude, whenever possible. As seen in this section, not providing this information amounts to considerable data over-reduction that leads to many subsequent issues^22^. Reporting effect estimates also helps safeguard against potentially spurious results. Signal changes in FMRI are relatively small and do not surpass 1-2%, except when simple sensory or motor conditions are contrasted with low-level baselines. In contrast, statistical values are dimensionless and do not directly provide information regarding effect magnitude. Indeed, the same statistic value may correspond, for example, to infinitely many possible pairs of mean and standard error (Fig. 4, right). A small *t*-statistic value could represent a small effect with a small standard error or a large effect with a large standard error – two scenarios with very different meanings. In addition, if, for example, a seemingly reasonable statistical value (e.g., *t*-value of 4.3) corresponds to an unphysiological 10% signal change, the conventional “statistic-only” reporting mechanism does not offer an easy avenue to identify and filter out such a spurious result. We note that another common practice of reporting a standardized “effect size” (e.g., Cohen’s *d*) shares the same problem of information loss due to (a) the unavailability of the physical scale and (b) the over-reduction from two values (effect estimate and uncertainty) to one. On the other hand, such a standardized metric, if desirable, could be easily derived from the effect estimate and its uncertainty.

The maturation of a field requires some extent of quantification and uncertainty assessment. However, currently most studies in neuroimaging only report binarized results at the level of qualitative assessment (e.g., positive or negative) with neither specifically quantified effect estimation nor uncertainty. Returning to the NARPS investigation, they performed a second meta-analysis solely based on statistic values. Under this approach without dichotomization, the findings across teams were substantially more consistent with one another, reaching a conclusion that was different from their first meta-analysis based on individual teams’ dichotomized reporting. These results are not only encouraging for the field of neuroimaging, but they also highlight the perils of the dichotomous approach. We conjecture that the meta-analysis results would have been further improved if both effect magnitude and uncertainty information had been incorporated in their meta-analyses. On the other hand, the conclusion bias would have been further exacerbated when results were binarized with “statistically nonsignificant” ones unreported and hidden.

The NARPS study, as a prototypical example in the field, highlights the importance of result reporting. For the primary study of interest, the analysis and modeling were set up for massively univariate analyses: inefficient modeling occurred because information was not shared globally across the brain, and adjustment for multiplicity was necessary. Results were required to be dichotomized in the form of “yes/no” decisions for a few specific regions; model comparison and validation were unnecessary. Only statistical evidence was presented in the results reporting, ignoring the informational context of effect magnitudes. The information loss due to these requirements, which mirror many conventional practices, can best be seen in the NARPS report by the contrasting conclusions these steps produce in a meta-analysis compared to a separate comparison done by looking at unthresholded statistics: the former “resulted in sizeable variation in the results of hypothesis tests”, while the latter “analyses of the underlying statistical parametric maps on which the hypothesis tests were based revealed greater consistency than would be expected from those inferences, and significant consensus in activated regions across teams was observed using meta-analysis.” That is, the consistency of results was noticeably greater just by loosening one of sources of information waste (dichotomizing). Similar to the demo example above in Fig. 5, it is likely that the finding of more consistency across teams represents reality more closely than the dichotomized version (which had undergone much greater information reduction before assessment). Much of the focus on the NARPS results has been on the “sizeable variation” of the dichotomized results; this has overshadowed the “significant consensus” that was present when the results were shown with less information loss. Due to the common practice of dichotomization and incomplete result reporting, meta-analysis in neuroimaging is largely limited to anatomical locations without regard to effect magnitude and is vulnerable to publication bias. Thus, information loss has a far-reaching impact on meta-analysis specifically but also on reproducibility in general.

To conclude this section, let us consider some of the issues discussed in the present and preceding sections. The common statistical practice in population-level analysis faces several challenges:

1. The principle of insufficient reason^9^, while reasonable in some statistical settings, in the case of FMRI disregards distributional information concerning effect magnitude across the brain (Fig. 3).
2. Hard thresholding carries with it a fair mount of arbitrariness and information waste.
3. The use of summary statistics alone to report results instead of a combination of effect estimate and uncertainty has detrimental impacts on study reproducibility (and makes spotting spurious results less straightforward).

Next, we describe how Bayesian multilevel modeling provides a paradigm to address these issues.

### 3.2 Bayesian multilevel modeling

We start with building up the structure of Bayesian multilevel modeling by first considering simple data *y_ij_*(*i* = 1, 2,…,*n*; *j* = 1, 2,…,*k*) from *n* subjects that are longitudinally measured under *k* time points with a predictor *x_ij_*, using the form *y_ij_* = *α_i_* + *β_i_x_ij_* + *ϵ_ij_* with intercept *α_i_*, slopes *β_i_*, and residuals *ϵ_ij_*. To appreciate the flexibility of the approach, this model is sometimes referred to as a “varying-intercept/varying-slope” model akin to those commonly adopted in a multilevel framework:

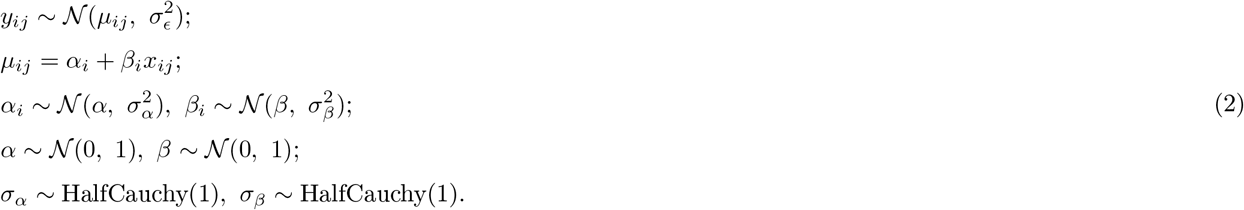

What makes the model “multilevel” is that it involves the hierarchical levels of subjects and time points. The notation *α_i_* indicates that each subject *i* has a unique intercept; likewise, *β_i_* shows that each subject *i* is given a unique slope. The first line specifies the *likelihood* or the distributional assumption for the data *y_ij_*. The expression for *μ_i_* specifies a linear relationship with a single predictor *x* (adding more predictors is straightforward). The third and fourth lines are *priors*: the varying intercepts follow a Gaussian distribution with a grand intercept *α* plus standard deviation *σ_α_*; likewise, the varying slopes follow a Gaussian distribution with a grand slope *β* plus standard deviation *σ_β_*. Importantly, the parameters of the prior distributions are learned from the data. Finally, the last two lines specify the *hyperpriors*, which can be conveniently weakly informative distributions for the means and standard deviations of the priors.

The above Bayesian multilevel modeling framework can be applied quite generally to any hierarchical structure. For example, meta-analysis is typically formulated under the conventional framework, as shown in the formula (1), through random-effects modeling. However, it can also be conceptualized as a Bayesian multilevel model as exemplified in Fig. 5. Even though the two approaches would often reach similar conclusions except for some degenerative cases^b^, the posterior distribution from Bayesian modeling provides richer information than a point estimate combined with a standard error. As illustrated in Fig. 5, we do not assume a uniform prior by adopting the principle of insufficient reason, nor do we adjust for multiple testing for individual studies as in the massively univariate approach. Rather, we regularize or apply partial pooling on the studies through weighting as shown in the formulation (1).

The Bayesian formulation (2) allows the modeler to flexibly estimate intercepts and slopes as a function of the hierarchical level of interest. Due to the impact of partial pooling across hierarchical levels, the Bayesian model tends to generate estimates that are more conservative and closer to the average effect within a given hierarchy than if each specific effect were estimated individually. Because of this conservative nature, the multilevel model aims to control for errors of incorrect magnitude and sign. Furthermore, adjustment for multiplicity is not needed^24^, especially since all the inferences are drawn from a single, overall posterior distribution of an integrative model.

In the past years, we have investigated how the framework can be effectively employed to analyze FMRI data at the region level^8,25,27^, as well as for matrix-based analysis including time series correlations among regions or white-matter properties^28^. The approach has also been applied effectively to other scenarios in neuroimaging^26,29–31^. Although at present the framework is computationally prohibitive at the whole-brain voxel level, we have also employed the technique at the voxel level within brain sectors, such as the insula. First, we present a region-level example that illustrates a recent application^25^ at the region level. The outcome in Fig. 6A shows the posterior distributions that characterize the probability of observing the region-level effects in a range given the data. For each region, the full posterior distribution conveys both the effect magnitude and its uncertainty, and the latter can be captured by the area under the curve to the right/left of zero. This posterior can be reported in full without dichotomization, as shown here. For example, the posterior probability that the effect was greater than zero in the left superior frontal gyrus (L SFG) was 0.92, which may be noteworthy in the research context in question. In particular, model fits can be qualitatively assessed by plotting predicted values against the raw data through posterior predictive checks (Fig. 6B) and quantitatively compared to alternative models using information criteria through leave-one-out cross-validation. By comparison, the model fit using the massively univariate approach was considerably poorer (Fig. 6B).

**Figure 6:**
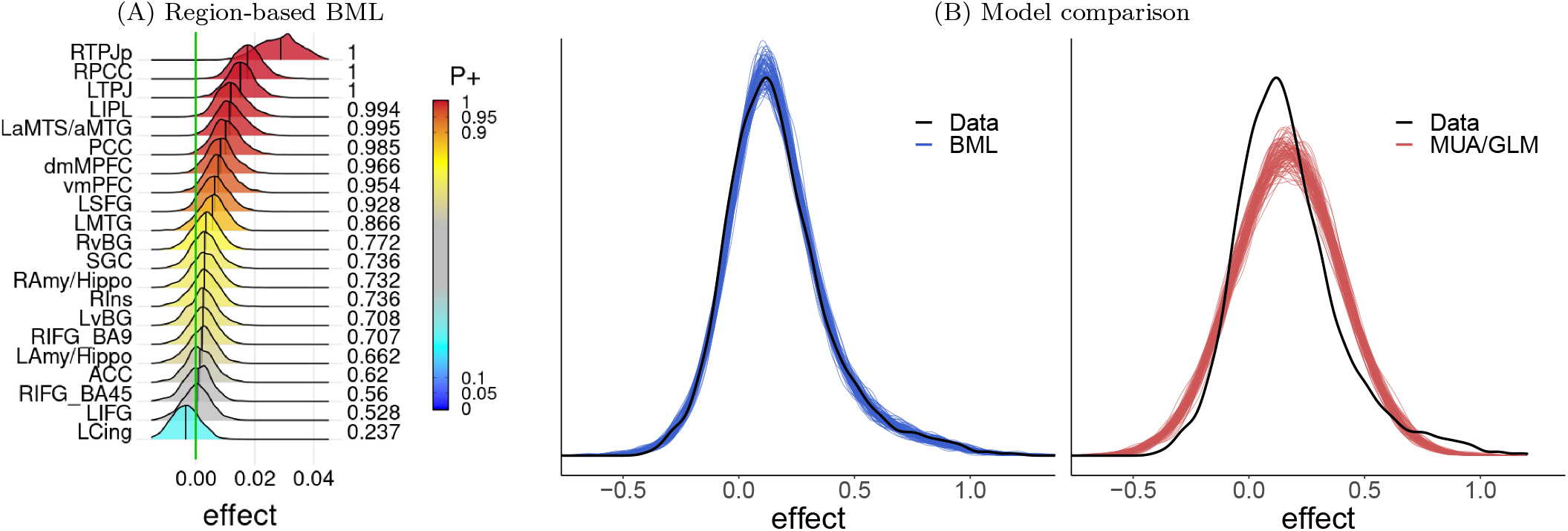
Bayesian multilevel (BML) modeling at the region level. (A) Population-level analysis was performed with an FMRI study of 124 subjects^25^. Colors represent values of *P*^+^: the posterior probability that the effect is positive. The analysis revealed that over one third of the regions exhibited considerable statistical evidence for a positive effect. In contrast, with massively univariate analysis, only two regions survived multiple testing adjustment^26^. (B) The BML performance is assessed and compared to the conventional approach. Posterior predictive checks visually compare model predictions against raw data. The BML model generated a better fit to the data compared to the general linear model (GLM) used in the massively univariate analysis (MUA).

The Bayesian multilevel approach can also be applied to voxel-level data within spatially delimited sectors. For instance, in a recent experiment, two separate groups of participants received mild electrical shocks^31^. In the *controllable* group, participants could control the termination of shocks by pressing a button; in the *uncontrollable* group, button pressing had no bearing on shock duration. The two groups were yoked so that, for a given participant, the exact timing of shock events in the controlled condition was replicated for a paired participant in the uncontrolled condition As in the standard FMRI approach, at the voxel level the effects (commonly denoted as *β* coefficients) of each participant were estimated based on a time series regression model. In the standard approach, one would proceed with voxelwise inferential tests (say, a *t*-test comparing the two groups) followed by a threshold adjustment based on spatial extent to control for multiple testing.

In contrast, the Bayesian multilevel approach specifies a single model, which combines all data according to natural hierarchical levels of the data. In this particular study, one natural hierarchy was that of participant pairs given the yoking of the experimental design. In addition, we focused on voxels within the insula, a cortical sector important for threat-related processing. However, the insula is a large and heterogeneous territory, with notable subdivisions that previously had been described functionally and anatomically. Accordingly, we subdivided the insula in each hemisphere into around 10 subregions, each of which with approximately 100 voxels. Thus, the subregions comprised another hierarchy. At the most basic level of the hierarchical structure, the unit was the voxel itself.

The following model was employed for the voxel-level data,

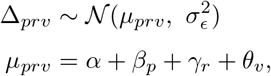

where the difference Δ_*prv*_ in FMRI responses to shock between a participant pair *p* in a voxel *v* belonging to region *r* is estimated at the subject level through time-series regression analysis and assumed to originate from a Gaussian distribution centered on *μ_prv_* with variance 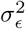. The second line specifies the response difference as a linear combination of an overall effect *α*, a contribution *β_p_* from participant pair *p*, a contribution *γ_r_* from region *r*, and a contribution *θ_v_* from voxel *v*. Importantly, the participant pairs, regions, and voxels are assumed to come from their respective (hypothetical) populations characterized by priors as in the model (2) (further specifications omitted here for brevity), and play a role equivalent to “random effects” in conventional linear mixed-effects models. Finally, for simplicity here we omitted several covariates that were included in the original analysis, including those related to individual differences in trait and state anxiety. Those covariates can be captured by slope parameters as in model (2), where it is possible to model them in terms of varying slopes (thus slopes can vary across regions, for example). This Bayesian machinery allows us to estimate the contributions of participant pairs, regions, and voxels based on the data, the likelihood, and the prior distributions. In the present study, our goal was to understand voxelwise effects (Fig. 7).

**Figure 7:**
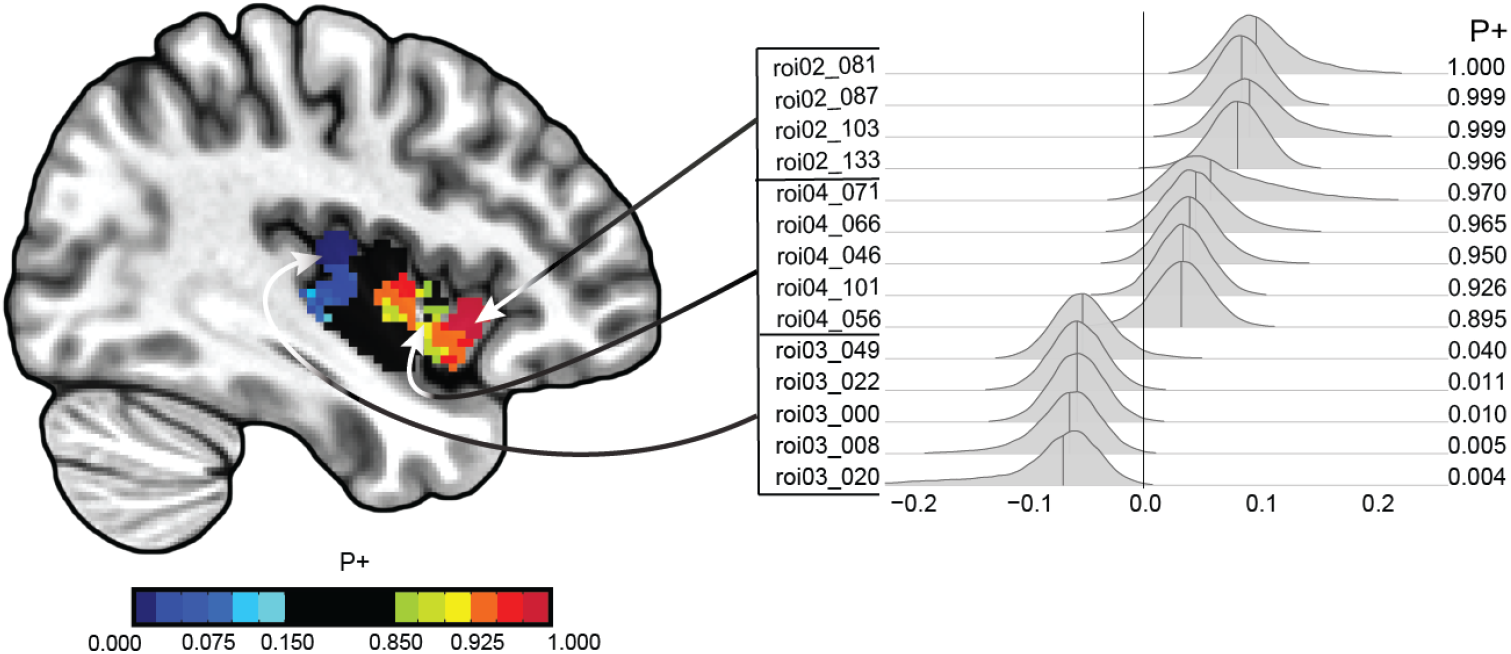
Bayesian multilevel voxelwise results. The right part of the figure illustrates posterior distributions of voxels from three subregions of the insula (voxels selected to illustrate some of the range of statistical evidence). Colors represent values of *P*+: the posterior probability that one condition (uncontrollable group) is greater than the other (controllable group). Values closer to 1 indicate stronger evidence that uncontrollable is greater than controllable, while values closer to 0 indicate the opposite (values computed based on the posterior distributions of the difference of the two conditions correspond to the tail areas of the posteriors). The computational time was about two weeks for this dataset with 126 subjects and approximately 1,000 voxels on a Linux server using 4 Markov chains.

To recapitulate, we note that the Bayesian approach can be adopted to achieve seven important goals.

1. *Handling multiplicity.* The Bayesian framework offers a potential avenue to addressing the problem of multiple testing that is so central to neuroimaging statistics. Because a *single* model is employed with information shared and regularized through partial pooling, all inferences are drawn from a single overall posterior distribution. Thus, information is more efficiently shared across multiple levels; no multiple testing adjustment is necessary^24^, avoiding excessive penalty due to information waste. In other words, instead of resorting to a post hoc adjustment for multiple testing under a modeling framework with an unrealistic assumption (e.g., uniform distribution), the Bayesian approach directly incorporates the interrelationship of the hierarchical structure as part of the modeling processing. We note that some statisticians have suggested other forms of adjustment based on decision theory.^2,32,33^
2. *No penalty against small regions.* Under massively univariate analysis, spatial extent is traded off against voxel-level statistical evidence in the process of adjusting for multiple testing. Thus, small regions are inherently placed in a disadvantageous position even if they have similar effect strength as larger ones. In contrast, under the Bayesian framework, each spatial unit is a priori assumed to be exchangeable from any other units. In other words, all units are a priori treated on an equal footing under one common prior distribution and are a posteri assessed on their own effect strength. As a result, small regions are not disadvantaged because of their anatomical size.^8^
3. *Insensitivity to data space.* Under the single integrative framework, the information is shared in a “melting pot” and calibrated. In other words, partial pooling plays a self-adaptive role of regularization, similar to the situation with the conventional methods such as ridge regression and LASSO. Under the single integrative framework, the information is shared in a “melting pot” and calibrated. In other words, partial pooling plays a self-adaptive role of regularization, similar to the situation with the conventional methods such as ridge regression and LASSO. Thus, the impact on the same spatial unit is relatively negligible even when the total amount of data changes (e.g., increasing the number of voxels or regions); as a result, a region would be assessed by its own “merits” of effect magnitude, but not its anatomical size.^8^
4. *Model quality control*. Model accuracy and adequacy can be assessed through posterior predictive checks and validations. For various reasons, model performance and comparisons are rarely cross-examined in neuroimaging. However, the modeling process should not be simply executed as an automatic pipeline without quality control. For example, a posterior predictive check allows one to examine the model adequacy or discrepancies through visually comparing the predictive distribution to the observed data. Cross-validation is another important technique under the Bayesian framework to gauge how closely a model, relative to potential candidates, predicts future data from the same data generating processes that produced the current data at hand. In general, the Bayesian approach welcomes an integrated view of the modeling workflow with an iterative process of model development and refinement^34^.
5. *Enhanced intepretability.* The Bayesian approach enhances the interpretability of analytical results. The posterior probability indicates the strength of the evidence associated with each effect estimate, conditioned on the data, model and priors. In the conventional null hypothesis framework, uncertainty is expressed in terms of standard error or confidence interval. Unfortunately, while mathematically precise, this information is very difficult to interpret in practice and easily misunderstood^35^. Notably, a confidence interval is “flat” in the sense that it does not carry distributional information; parameter values in the middle of a confidence interval are not necessarily more or less likely than those close to the end points of the interval, for example (e.g., Fig. 5A,C). In contrast, the posterior distribution provides quantitative information about the probability of ranges of values, such as the parameter being positive, negative, or within a particular interval. Unlike the conventional notion of “confidence interval”, parameter values surrounding the peak of the posterior distribution are more likely than those at the extremes (Fig. 5B).
6. *Error controllability.* Instead of the false positive and false negative errors associated with the conventional null hypothesis framework, Bayesian multilevel modeling can be used to control two different errors: *type M* (over- or under-estimation of effect magnitude) and *type S* (incorrect sign)^36^. For example, effect estimates under the massive univariate modeling framework tend to be exaggerated (Jensen’s inequality), leading to type M error. In contrast, shrinking the effect estimates under the Bayesian multilevel framework provides an effective way to regularize and counteract the exaggeration ^14^.
7. *Extended modeling capabilities.* The Bayesian framework is advantageous and flexible in handling complex data structures that can be challenging for the conventional framework. Consistent with the central limit theorem, many types of data, including the effect estimates in neuroimaging, tend to have a density of roughly Gaussian characteristics with a bell-shaped distribution exhibiting a single peak and near symmetry. Based on the maximum entropy principle, the most conservative distribution is the Gaussian if the data have a finite variance^37^. Thus, for the same reasons that subjects in neuroimaging are routinely treated as random samples from a hypothetical pool of a Gaussian distribution, we can effectively model the effect distribution across space as a Gaussian, rather than adopting the stance of “full ignorance”. However, exceptions do occur when the data do not follow a bell-shaped distribution due to outliers or skewness. A conventional approach is to set hard bounds (e.g., the rule of three standard deviations), constraining the data to a predetermined interval in order to exclude outliers. Such a brute force approach is arbitrary and unprincipled to some extent. In contrast, outliers or skewed data can be accommodated in a principled manner with the utilization of non-Gaussian distributions (e.g., Student t-distribution with an adaptive number of degrees of freedom, Lambert W transforms) for data variability. Another benefit of Bayesian modeling is the convenience of incorporating the uncertainty information (e.g., measurement errors) for both response and explanatory variables that may improve modeling accuracy and accommodate data asymmetry due to outliers or data skewness.

Theoretically the Bayesian multilevel framework can incorporate any number (large or small) of spatial units, voxels or regions, into one unified model. Numerical considerations aside, such a Bayesian model is essentially the same as the traditional linear mixed-effect formulation both in conceptual viewpoint and in symbolic model expression. In addition, the crucial aspect of the hierarchical framework lies in the assumption of, for example, a Gaussian distribution for the variability across spatial units. It is this distribution assumption that plays the role of information sharing across space through global calibration or shrinkage. If the prior is relaxed from a Gaussian distribution to a trivial case of uniform distribution, then no information is shared across the spatial units per the principle of indifference that assigns epistemic probabilities^9^. In other words, in the absence of available evidence, one could adopt a uniform distribution across space; thus, the hierarchical model would simply reduce to a special case: namely, the conventional massively univariate model. However, as discussed here, it is generally more reasonable as well as more informationally efficient to adopt a hierarchical model with the assumption of an approximately bell-shaped, rather than uniform, distribution for cross-spatial variability, as evidenced in the empirical data of Fig. 3B.

The adoption of Bayesian multilevel modeling here is intended to specifically address information waste through three issues at the population level: mischaracterization of data hierarchies, dichotomization and data over-reduction. There has been a rich literature of Bayesian applications in neuroimaging that focus on various aspects of modeling. The wide range of topics include: using Bayesian modeling as an alternative at the subject level (e.g., temporal structure^38^, spatio-temporal modeling^39^, complex-valued FMRI^40^, hemodynamic response estimation and connectivity^41^, empirical Bayes for spatio-temporal modeling^42–48^), adopting empirical Bayes to resolve the unstable cross-subject variability (e.g., when the number of subjects is 40 or less) by sharing data variability across neighboring voxels^49^, leveraging between anatomical data with a high resolution and those with a low resolution^50^, handling measurement errors^51^. Some of these methods have adopted a similar concept of partial pooling for time series regression at the subject level^42,44,45,48^ or for locally regularizing cross-subject variability at the population level^49^. However, it is beyond the scope and space in this commentary to provide an exhaustive and detailed coverage for these topics that are tangential to our focus of addressing the issue of information loss at the population level.

### 3.3 Neuroimaging without *p*-value thresholds?

Let us consider the issue of probability thresholding, regardless of the modeling framework, in further detail. Dichotomization is essential to statistically-based decision making. As noted above, it provides a way to filter a lot of information and to present results in a highly digestible form: binary ON/OFF output. For example, based on the available data, should a certain vaccine be administered to prevent Covid-19? In such cases, a binary decision must be adopted, and decision theory, which incorporates the costs of both false positives and false negatives, can be used. Here, we entertain a seemingly radical proposal: What would be lost in neuroimaging if hard thresholds were abandoned? It could be argued that this would lead to an explosion of unsubstantiated findings that would flood the literature. We believe this is unlikely to occur. Scientists are interested in finding the probability of seeing the effect conditioned on the data at hand (“what happened”), rather than the *p*-value (“what might have happened” or the probability of seeing the data or more extreme scenarios conditioned on the null effect). The absence of a hard threshold does not entail that “anything goes”; rather, it encourages substituting a mechanical rule by careful justification of the noteworthiness of the findings in a larger context.

Consider the controllability study discussed above. In additional analyses at the level of brain regions, we found very strong evidence (*P*+ = 0.99) for a controllability effect in the bed nucleus of the stria terminalis, a structure that plays an important role in the processing of threat. This region and the central nucleus of the amygdala are frequently conceptualized as part of a functional system called the “extended amygdala”. Accordingly, we found it important to emphasize that there was also some evidence (*P*+ = 0.90) for a controllability effect in the left central amygdala. Although the central amygdala did not meet typical statistical cut-offs, we believe that the finding is noteworthy in the larger context of threat-related processing. This is particularly the case because reporting the central amygdala effect can be informative when integrating it with other studies to perform meta-analysis, as discussed in Section 3.1. Note that by providing the information about the central amygdala, readers are free to interpret the findings in whatever way they prefer; they may agree with our interpretation (that there is some evidence for an effect in this region), or consider the evidence “just too weak”. This is not a problem in our view; rather, it is a feature of the approach we advocate for.

A more flexible approach both in terms of statistical modeling and in terms of result reporting is potentially beneficial. At the heart of the scientific enterprise is rigor. In experimental research, typically this translates into testing patterns in data in terms of null hypotheses and a *p*-value threshold of 0.05. On the surface, the precise cutoff provides an objective standard that reviewers and journal editors can abide by. On the other hand, the use of a strict threshold comes with its own consequences. In most research areas, including neuroimaging, data are notoriously variable and not readily accommodated by simple models^37^. In this context, is it really essential to treat a cluster size of, say, 54 voxels as qualitatively different from one with 50 voxels? As models by definition have limitations, we believe that dichotomization, as illustrated by the example in Fig. 5, is unproductive.

In light of these considerations, we propose a more “holistic” approach that integrates both *quantitative* and *qualitative* dimensions. A recent investigation through Bayesian multilevel modeling indicates that full result reporting including visualization can effectively replace dichotomous thinking^52^. For results based on the conventional framework, we suggest a general *highlight but not hide* approach. Instead of applying a threshold that excludes results that do not cross it, one can show all (or most) results while highlighting or differentiating different levels of statistical evidence^53^ (Fig. 1F). Similarly, tables can include regions with a broad spectrum of statistical evidence, together with both their effect magnitudes and uncertainties. Overall, probability values, including the conventional *p*-value based on null-hypothesis testing, play a role as a piece of information, rather than serving a gate-keeping function. In addition, we encourage a mindset of “accepting uncertainty and embracing variation”^54^ in the results of any particular study.

### 3.4 Modeling trial-by-trial variability

In this section, we further illustrate the potential of using the Bayesian multilevel approach to build integrative analysis frameworks. In FMRI experiments, the interest is usually on various comparisons at the condition level. As condition-level effects exhibit considerable variability, researchers rely on multiple trial repetitions of a given condition to estimate the response via a process that essentially amounts to averaging. In this manner, trial-by-trial variability is often treated as noise under the assumption that a “true” response exists, and deviations from it constitute random variability originating from the measurement itself or from neuronal/hemodynamic sources.

However, neglecting trial-by-trial variability means that trial-level effects are considered as “fixed” in the fixed vs. random effects terminology, as opposed to participants, which are treated as random and sampled from a hypothetical population. Technically, this means that researchers cannot generalize beyond the specific stimuli employed in the experiment (say, the 20 faces used from a given dataset), as recognized several decades ago^55,56^. By modeling trials as varying instantiations of an idealized condition, a study can generalize the results to trials beyond the confine of those employed in the experiment^57,58^. Consider a segment of a simple experiment presenting five faces. In the standard approach, the time series is modeled with a single regressor that takes into account all face instances (Fig. 8a-b). The fit does a reasonable job at capturing the mean response; however, it is clearly poor in explaining the fluctuations at the trial level (Fig. 8c). Whereas traditional models in neuroimaging ignore this variability across trials, we propose to explicitly account for it in the underlying statistical model ^57,58^.

**Figure 8:**
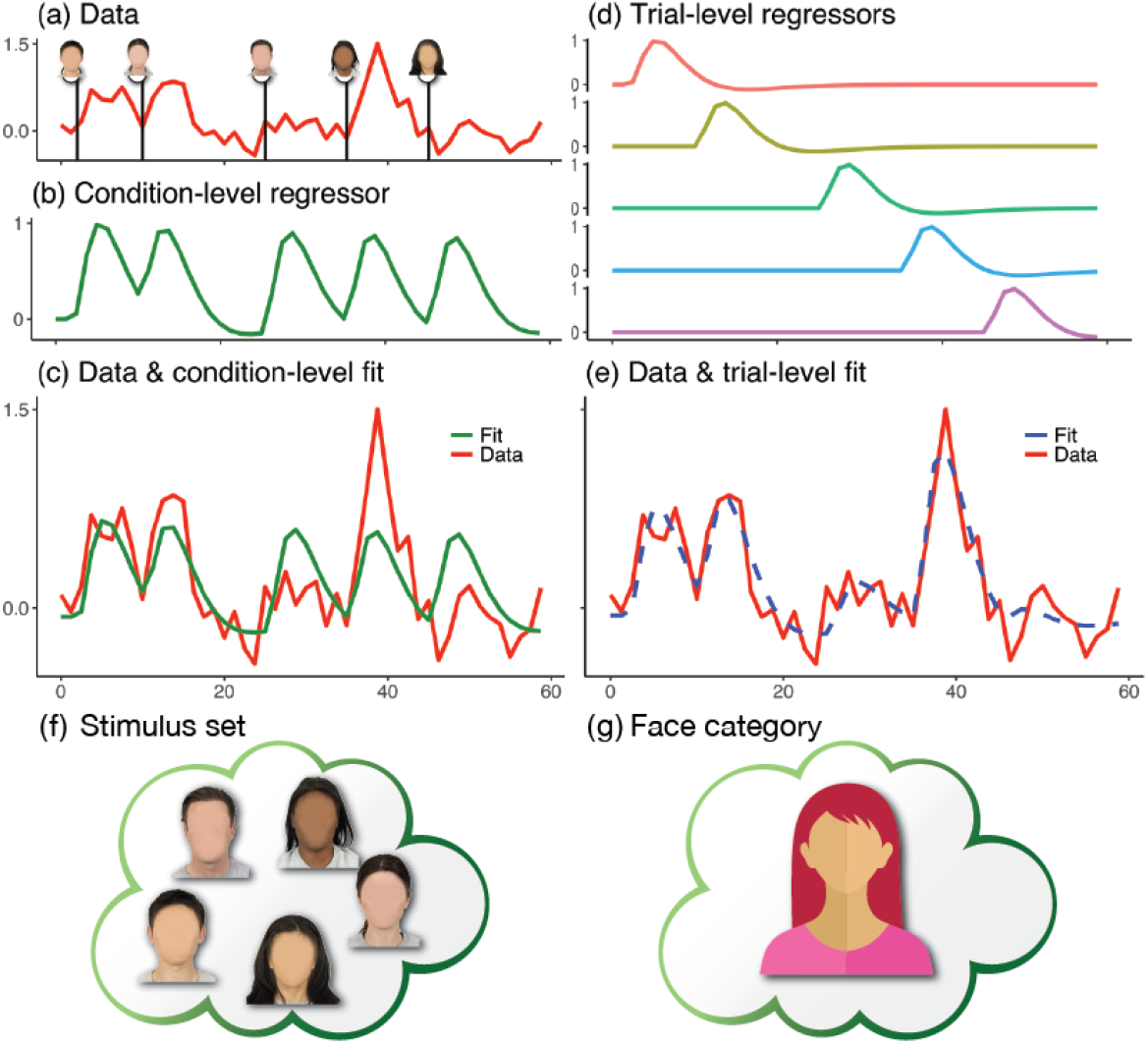
Time series modeling and trialbased analysis. Consider an experiment with five face stimuli. (a) Hypothetical times series. (b) The conventional modeling approach assumes that all stimuli produce the same response, so one regressor is employed. (c) Condition-level effect (e.g., in percent signal change) is estimated through the regressor fit (green). (d-e) Trial-based modeling employs a separate regressor per stimulus, improving the fit (dashed blue). (f-g) Technically, the condition-level modeling allows inferences to be made at the level of the specific stimulus set utilized, whereas the trial-based approach allows generalization to a face category.

The Bayesian multilevel framework can directly be used to account for trial-level effects. Specifically, at the subject level, we construct regressors for individual trials as in Fig. 8d. In a recent study, we explored a series of population-level models of trial-by-trial variability for FMRI data^58^, and indeed observed considerable cross-trial variation and notable inferential differences when trials were explicitly modeled. For example, as the experiment included a task involving negative or neutral faces, we were interested in amygdala responses, but our interest extended to a trial phase only containing cues indicating whether the trial was rewarded or not (in reward trials, participants received extra cash for correct and timely responses). Fig. 9 shows that trial-level modeling provided considerably stronger evidence for an effect of reward in the amygdala compared to the conventional condition-level modeling.

**Figure 9:**
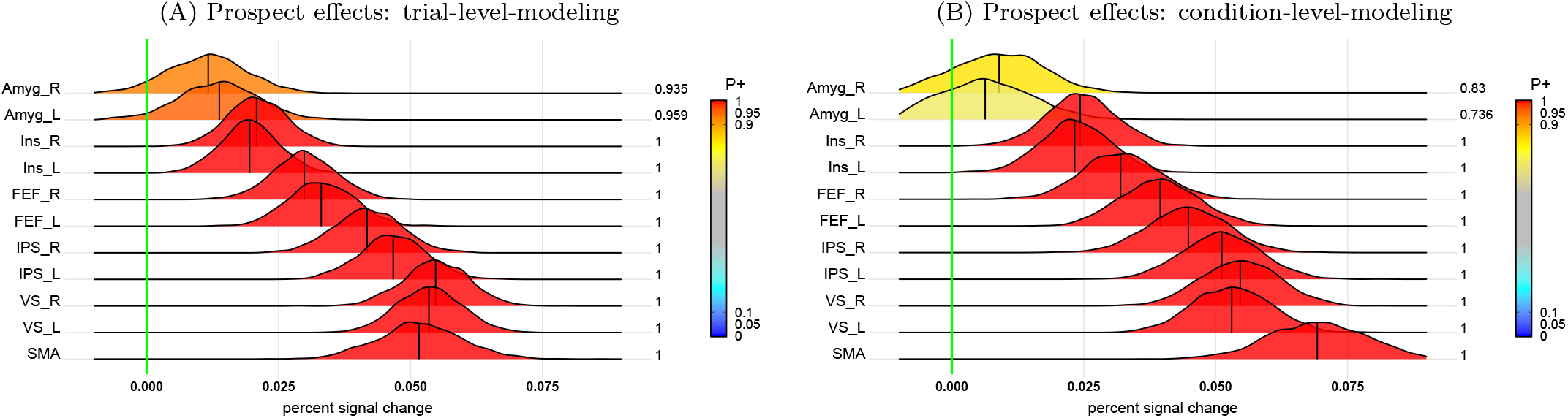
Trial-level versus condition-level modeling. Posterior distributions for the effect of reward (vs. control) cues for each region of interest. Although the two approaches provided comparable results, trial-level modeling (A) showed stronger evidence for left and right amygdala than the condition-level counterpart (B).

Trial-level modeling also improves the estimation of test-retest reliability (i.e., the degree of agreement or consistency of subject-level differences between two or more repeated measurements). Recent reports have suggested that the test-retest reliability for psychometric^59^ and neuroimaging^60^ data is rather low when evaluated via the conventional intraclass correlation coefficient. The low reliability of effects with robust population-level effects (e.g., Stroop and Flanker tasks) was particularly worrisome in the context of individual-differences research. In a recent study, we developed a multilevel modeling framework that takes into account the data hierarchies down to the trial level, providing a test-retest reliability formulation that is disentangled from trial-level variability^27^. As a result, the trial-level modeling approach revealed the attenuation when the conventional intraclass correlation coefficient is adopted, and improved the accuracy of reliability estimation in assessing individual differences.

Two complex issues about trial-level modeling are experimental design and trial sample size. When the effect at each trial is separately characterized in the model at the subject level, high correlations or multicollinearity may arise among the regressors. To avoid such potential issues, trial sequence and timing can be randomized in the experimental design. As shown in a few recent studies^27,58,61^, even fast event-related experiments with a short inter-trial interval can be carefully designed so that trial-level effects can be captured. Nevertheless, detailed attention is still needed in processing and quality control, because unstable effect estimates, outliers, and skewed distributions may still occur due to high collinearity among neighboring trials or head motion. Our recent investigations^27,58,61^ provide some solutions to handle such difficult situations. Furthermore, even though trial sample size is largely chosen as a convenient or conventional number with which the subject would be able to endure during the scanning session, our recent investigation^61^ indicates that it has nearly the same impact as subject sample size on statistical efficiency.

## 4 Discussion

Neuroimaging research is challenging, not least because data analysis includes several interdependent steps of processing and modeling. Data from tens of thousands of spatial units are acquired as a function of time for one or multiple subject groups and for several experimental conditions with trials repeated many times per condition, typically across multiple data acquisition runs. Given the challenges any one research team would face to analyze this type of data, developers have designed software packages that enormously lower the barrier to entry to investigators. Indeed, statistical development for FMRI analysis has proceeded vigorously since the early 1990s. Among the greatest challenges has been the issue of multiple testing, with the dream of “whole-brain noninvasive” imaging coming at a severe cost inferentially. Since the beginning, experimenters have been admonished that without “strict enough” procedures, the “false positive rate” would be prohibitively high. Accordingly, considerable research has been devoted to to improving inferential efficiency.

### 4.1 Conditionality of statistical information

Contrary to common practice, the strength and accuracy of statistical evidence are not as informative as usually perceived. According to the central limit theorem, given a large enough sample size (e.g., big data initiatives), statistical evidence may reach as strong as any designated level unless the associated effect is absolutely zero. For example, suppose that the contrast between positive and negative valences in a brain region follows a Gaussian distribution 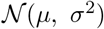 with *μ* = 0.2, *σ* = 0.5 (in units of percent signal change). With a sample size of 40 or 1,000 subjects, one could condense and reduce the data information, as typically done in the literature, to a single number, such as *p* ≈ 0.01 or *p* ≈ 1.0 × 10^−22^, respectively. It is difficult to properly digest the information from these two statistical values and to assess as to how much more information is contained in the latter than the former. In contrast, much more revealing information could be attained if their posterior distributions or corresponding 95% uncertainty intervals of BOLD effect (e.g., (0.04, 0.36) and (0.17, 0.23)) were provided.

Statistical interpretation should be properly framed and contextualized. In the aforementioned example, an uncertainty interval only characterizes the population average *μ*, which is largely a theoretical or abstract construct; thus, one should not lose the sight when gauging a particular subject’s effect, which could vary in a much larger range (cf., 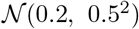). In addition, any statistical inference is conditional on the adopted modeling framework, the underlying assumptions, and the representativeness of the sampled data. For instance, all statistical models are to some extent an idealization or approximation of the reality; pragmatically, analytical decisions require scientific evaluation, including specific aspects of data processing (amount of spatial smoothing, choice of data included or excluded, model validation, etc.) and uncertainty assignment through a probability distribution. Therefore, estimation accuracy, statistical evidence, and probabilistic reasoning may change if, for example, confounding variables, interaction effects, and nonlinearity are incorporated, when a different distribution is assumed, or whether some extent of regularization is applied.

### 4.2 Applicability of Bayesian multilevel framework

Bayesian modeling in general has several advantages that could substantially benefit the neuroimaging community, but some barriers at present hinder its wide adoption. We list here a few directly relevant, but not exhaustive, advantages:

1. natural, intuitive and straightforward interpretation based on probabilistic inference,
2. the convenience of incorporating prior information (e.g., distribution across spatial units),
3. the capability of accurately capturing the data generative mechanism,
4. efficient handling of multiplicity through partial pooling,
5. no penalty against regions because of their small anatomical size,
6. strong power of numerical solutions through Monte Carlo simulations,
7. full result reporting that contains both effect magnitude and uncertainty,
8. built-in model comparisons and quality check.

In contrast, at least three negative aspects prevent the Bayesian framework from its wide application: two of them are educational and the third practical. First, in most classrooms the teaching content of Bayesian modeling is sparse and generally quite outdated. As a result, most investigators are not well-versed to quickly adopt the concept and structure of the modeling framework. Second, historically there is a negative impression of “subjectivity” associated with the notion of priors. Third, computational hurdles have slowed the uptake of Bayesian applications until the numerical breakthrough of Markov chain Monte Carlo simulations.

Despite these hurdles, we are optimistic on the expanding potential of Bayesian modeling, as each barrier has been decreasing over time. Firstly, there is more awareness and educational momentum about these methods, increasing their popularity, application and development. For instance, a Bayesian multilevel model usually has a counterpart in the conventional linear mixed-effect formulation and contains the latter as a special case. Thus, the realization of this fact may help dissolve some of the difficulties for those who are accustomed to the conventional paradigm. Secondly, while Bayesian modeling does need to choose priors, one must remember that conventional massively univariate approach also makes an implicit prior assumption (a uniform distribution across space), leading to information waste. One must also remember that the goal of modeling is to closely characterize the data hierarchies: Bayesian modeling is more powerful in the sense that its performance and the adoption of priors can be rigorously evaluated via graphical tools such as posterior predictive checks and cross-validations (Fig. 6). Finally, in terms of current practical limitations, we are optimistic that, as is often the case with the computational costs of new methods, further development and innovations will increase speed and processing time; this may be naive, but it certainly seems reasonable given trends in neuroimaging data analysis as a whole over the past couple of decades.

Hierarchical modeling provides a suitable platform for closely characterizing the data generative mechanism, and Bayesian multilevel modeling offers an important numerical machinery in making statistical inferences. Due to the complex and intertwining data structures in neuroimaging that involves many levels such as time series, trials, conditions, subjects, groups and brain regions, it is pivotal to adopt a holistic perspective that reflects as close as possible to the sources of data variability. It is also this hierarchical consideration that motivates us to propose the incorporation of spatial units as part of the modeling process as opposed to a post hoc compensation for multiple testing under the conventional massively univariate framework. Some of the hierarchical schemata are conceptually equivalent to various regularization methods (e.g., ridge, LASSO); they can be also formulated under the conventional mixed-effects paradigm with, for example, the spatial units playing the role of “random effects”. Although computationally affordable, a linear mixed-effects model could only allow one to make inferences at the population level, but not for spatial units as individual random effects. In contrast, Monte Carlo simulations, despite the relatively high numerical cost, enable the Bayesian framework to accommodate a wide range of modeling capability and inferential power. We offer three programs of Bayesian multilevel modeling in neuroimaging as part of the AFNI suite for public use: **RBA**^25^ and **MBA**^28^ for region- and matrix-based analysis, and **TRR**^27^ for test-retest reliability estimation.

### 4.3 Analytical level: voxel versus region

The choice of analysis level – voxel or region – deserves some elaboration and discourse. Both are commonly used across the neuroimaging field, and the choice involves some trade-offs in processing and modeling at the population level; it can even affect final interpretation (though, some differences are not as large as they might appear).

Modeling at the voxel level has the benefits of relatively high spatial resolution and independence of a choice of region definition. However, as the inferential focus is usually on the cluster – not voxel – level, one thorny issue is the lack of spatial specificity that plagues the massively univariate framework. A post hoc solution^62^ has been proposed to address and improve the issue of lacking spatial specificity; this provides an interesting approach, which could also be integrated with other issues raised here (e.g., lack of region-level uncertainty, information loss due to the absence of global calibration). Nevertheless, the actual spatial resolution of signals is not actually the acquired voxel size (nor any upsampled final size). There is inherent smoothness in the acquired data and more is added by blurring during preprocessing; this blurring decreases spatial resolution by spreading information across anatomical structures, which makes final interpretation more difficult and less spatially specific. The initial spatial specificity is further lessened by the post hoc adjustment for multiplicity when modeling at the voxel level through massively univariate analysis, which commonly relies on spatial clustering. In the end, individual voxels are not interpretable, and only surviving clusters are.

Additionally, voxelwise analyses are not independent of brain regions: the location of voxels within anatomical structures still matters in multiple ways. In practice, most adjustments for multiplicity penalize small regions. Importantly, even voxelwise studies aim to produce and make inferences at the region level. The typical research hypothesis is framed in terms of regions of interest, such as, “Based on previous literature, we hypothesize that regions X, Y and Z will show…” A recent example, the NARPS study described above, asked teams to report yes/no findings about certain regions of interest, which were then combined for a meta-analysis (although a specific atlas was not specified; researchers had to choose their own definitions).

Moreover, voxelwise analyses have been susceptible to problematic or inconsistent results reporting. While these issues are not strictly inherent to the approach, their prevalence makes it difficult to disentangle. Often, there is overfitting at the voxel level, leading simultaneously to exaggerated estimates and poor prediction accuracy. Furthermore, with dichotomization through clustering, interpretational difficulties can arise. Typically, each cluster is not localized within a single brain region – should all the regions be reported, or just those with large overlap (how to define “large”?)? Many researchers just report the location of a single voxel with “peak” statistical values, in order to summarize the results for the entire cluster. This is statistically inconsistent with the thresholding procedures, and fundamentally a large source of information waste.

Region-based studies have some drawbacks as well. For example, there is a semi-arbitrariness of parcellation selection, which will affect results. Furthermore, not all structures of interest exist in available parcellations (though the number available will surely only increase over time). With multiple regions analyzed through conventional approaches, the penalty for multiplicity is often quite high, to the point of being unacceptable and/or unrealistic. In some cases, subregion differentiation can be important (e.g., localizing focal lesions), but these tend to be clinical cases and not often occurring in population-level studies.

However, there are several useful properties of region-based analyses. Firstly, there is a meaningful specificity to results: in most brain studies, including NARPS, research hypotheses are based on regions, and in this case the list of relevant regions is clearly defined from the start and consistent with the study design. Blurring is not included in the preprocessing, so signals are not spread widely across regions. Having a smaller number of spatial units means analyses take less computational time and cost, and also that it is easier to avoid dichotomization. Practically, modeling at the region level opens the door to capture the hierarchical structures and to improve model efficiency. Finally, all regions – big or small – are treated equally.

All things considered, region-based and voxelwise analyses at the population level share several commonalities. Each relies heavily on alignment, even if in different ways. By the time modeling is complete, the spatial resolution of the two approaches is likely not very different, depending on the parcellation. In fact, one can note that recent parcellations have created more and more regions in a standard template brain (greater than several hundred regions, in some cases), so that their spatial resolution has become finer. Indeed, at this level voxels are then just the limiting case of finer parcellations, particularly once blurring has been accounted for. In both cases, what remains most important is to utilize a reasonable modeling framework and statistical practices with either methodology, reducing inconsistency and information waste to the greatest extent possible.

### 4.4 Maintaining enough information in result reporting

The conventional massively univariate analysis as an exploratory tool can benefit from our discussion here regarding the principles and rationales underlying the Bayesian framework. Because of the computational burden, currently the Bayesian multilevel model can only afford to handle up to a few thousand spatial units (e.g., regions or voxels); thus, exploratory analysis through whole-brain voxel-level is presently beyond the reach of Bayesian modeling. However, future methodological developments and computational breakthroughs will surely continue to reduce, if not dissolve, this computational barrier^63,64^. Nevertheless, we believe that the hierarchical perspective helps reveal the information loss associated with two aspects of the conventional modeling approach: the implicit assumption of uniform distribution and the artificial dichotomization required in handling multiplicity. In fact, these two aspects are two sides of the same coin: the conventional modeling methodology focuses only on local relatedness among neighboring spatial units, but ignores the global information shared across the whole brain. Consequently, the various approaches of adjustment for multiple testing adopted in the field may lead to excessive penalties and overconservative inferences. For these considerations, when voxelwise analysis is performed under the conventional massively univariate framework, we believe that the Bayesian multilevel framework lends an important perspective: a threshold or a set of spatial blobs purely based on statistical evidence is only suggestive but not rigid. To avoid further information waste, any statistical evidence should be viewed – regardless of the adopted framework – as intrinsically embedded with some underlying and implicit assumptions; it should be considered as a continuum both in result reporting as well as during the research reviewing process.

Here, we have addressed a few issues within conventional neuroimaging analysis pipelines: in the process of breaking down raw data and turning it into understandable results, we do not focus on boiling everything down to a small number of ON locations (in a sea of OFF background) at a given statistical significance level. We have shown the many ways that this can be considered an “overdigestible” result: a lot of useful information has been sacriñced (results at subthreshold locations that might still be informative, and separate effect estimates with uncertainty measures) for not much gain. Additionally, we have demonstrated that the conventional modeling approach is inefficient and wastes data, even before getting to questions of dichotomization: the implicit assumption of uniform distribution is far from approximating any realistic brain effect, and the *p*-values only provide limited information about how unlikely the current data or more extreme observations would be if a null effect *were* true, rather than the probability of research hypothesis being true *given* the data present.

Instead, we have proposed a small but important improvement to standard neuroimaging pipelines with an approach that aims to make more efficient use of the initial data, and that also has positive side effects for scientific inquiries. A schematic of this approach is shown in Fig. 10, in direct comparison with the traditional approach in terms of information loss and digestibility. Firstly, the Bayesian multilevel modeling approach replaces the massively univariate analysis and the principle of insufficient reason with a single integrative model and removes any later need for multiple testing adjustment. One benefit of this approach is now obtaining an overall posterior distribution for all model parameters, which provides a great deal of useful information about the estimate uncertainty as well as the overall model fit. This procedure also employs partial pooling across spatial units, so that the effect estimates are regularized to avoid potential overfitting.

**Figure 10:**
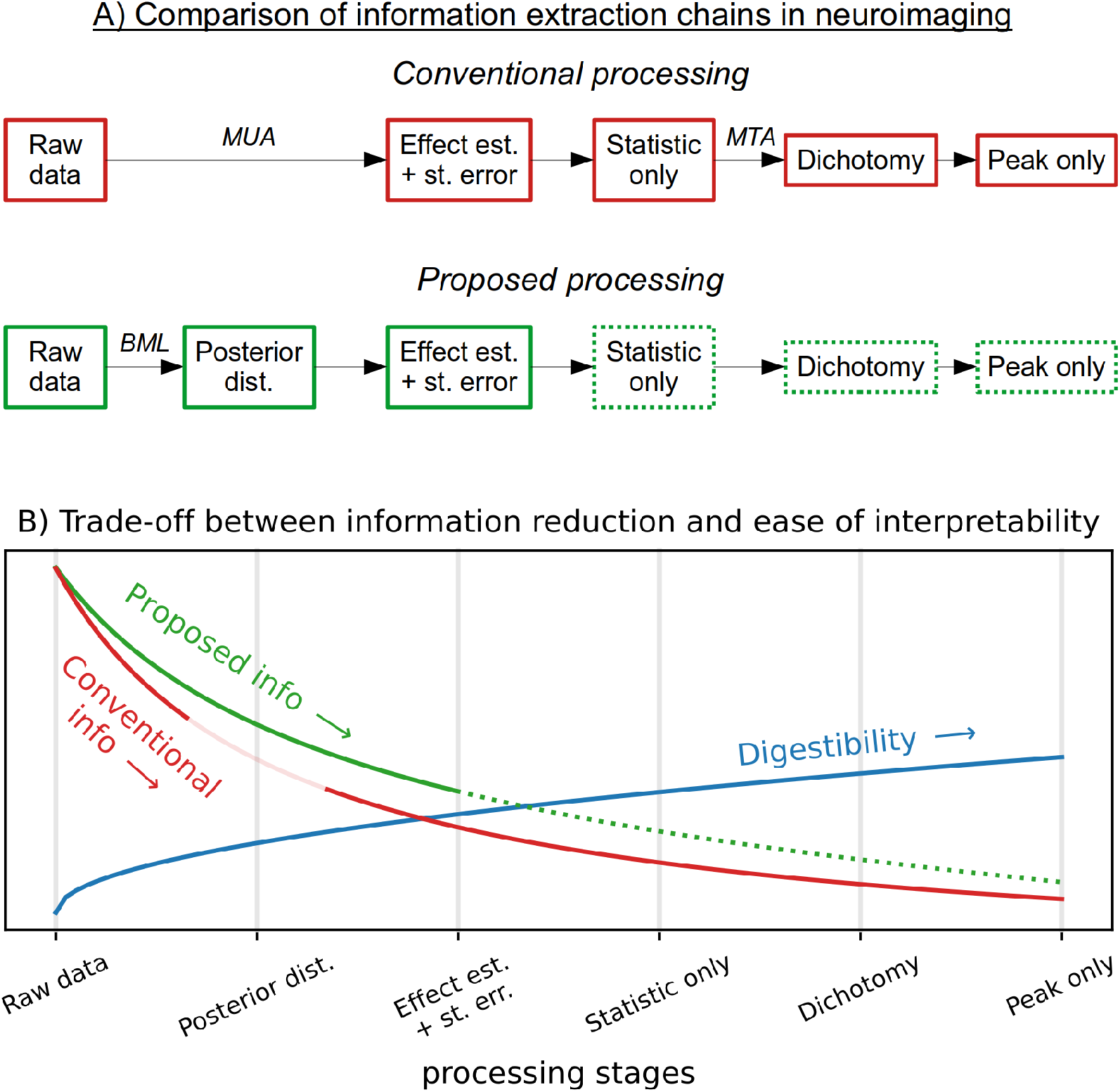
Comparison of FMRI information extraction for conventional and proposed Bayesian multilevel (BML) approaches (cf. Fig. 2). (A) The two approaches run parallel, but in the “proposed” first step BML puts data into a single model (removing the need for multiple testing adjustment later), and the information is partially pooled and shared across space. (B) The proposed multilevel framework produces an intermediate output of posterior distributions (lacking in the conventional approach); these carry rich information about parameter and model fitting. The information pooling also produces effect estimates that retain more information and avoid potential overfitting. This information advantage over the conventional method carries on to later stages. Thus, while the “digestibility” of results increases similarly at each stage, the drop-off in information content is slower in the proposed approach. The dotted part of the proposed steps reflects that we strongly suggest not including the steps that many traditional approaches at present perform, due to the wasteful information loss incurred.

Our concrete suggestions in result reporting are as follows. For whole-brain voxelwise analysis under the conventional framework, we recommend that one adopt a “highlight but not hide” approach. Specifically, it is preferable to highlight brain regions with some extent of statistical evidence (e.g., a cluster threshold of 10 voxels at the voxelwise *p*-value of 0.05) while gradually fading away for the rest (Fig. 1F), instead of the conventional dichotomization methodology (Fig. 1D). Region-level results under the conventional framework can be reported in a table that should include *both* the estimation of effect magnitude *and* the corresponding uncertainty (standard error or uncertainty interval). For region-based analysis under the Bayesian framework, we suggest, if space permits, the adoption of a more informative presentation than a table by showing each full posterior distribution as illustrated in Figs. 6A, 7, and 9.

We note that the information contained in the Bayesian results is much richer and more straightforward in result interpretation. For example, the posterior distributions in Figs. 6A, 7, and 9 show the full range of effect estimates and their uncertainty. In addition, each posterior distribution directly reveals the probability of seeing the effect in any range (e.g., being positive) conditional on the current data. As a comparison, even if the analysis under the massively univariate framework is performed at the voxel level, the significance level (e.g., 0.05) adjusted for multiple testing can only be applied to a whole set of spatial blobs, not to individual voxels, due to dichotomization; thus, it would be difficult to attach some sense of uncertainty for each spatial blob.

Our assessment and recommendation regarding modeling and result communication are summarized in Fig. 10. As of 2021, investigators have at their disposal a vast array of tools for the statistical analysis of FMRI data. The majority of them maintain a traditional focus on the conventional way of thinking of inferences in terms of “true” and “false” effects. In the present paper, we discussed several problems with applying standard null hypothesis significance testing to FMRI data. We favor a view of neuroimaging effects in terms of a continuum of statistical evidence, with a large number of small effects dominating, instead of islands of strong/true effects that should be discerned from false positives. We propose that Bayesian multilevel modeling has considerable potential in complementing, if not improving, statistical practices in the field, one that emphasizes effect estimation rather than statistical dichotomization, with the goal of “seeing the forest for the trees” and improving the quality and reproducibility of research in the field.

In neuroimaging, research groups acquire different sized datasets with different sample sizes and paradigms varying to some degree. With various preprocessing and modeling approaches available in the community, some extent of result variation is expected and unavoidable. All these factors contribute to an expected variability in reported results, and it need not be considered inherently problematic. To accurately combine multiple studies and determine the levels of variability present, one would need to make a model using their *un*thresholded results, and preferable both their effect estimates and uncertainty information. Otherwise, small outcome differences can appear to be much larger, when passed through the dichotomization sieve. Thus, it is the result presentation (e.g. highlight but not hide, show effect magnitude instead of statistical evidence only, revealing model details, etc.) that would conduce to the convergence of a specific research hypothesis across teams. We believe that the abandoning of result dichotomization is one small step toward reducing variability due to artificial thresholding. We agree with NARPS’s suggestion of encouraging original statistical results being submitted to a public site. However, more improvements would be needed. For example, such public results at present are still restricted to statistical evidence without the availability of effect magnitude information. Furthermore, proper presentations in publications remain a crucial interface for direct scientific communication and exchange. Therefore, in repositories such as NeuroVault^65^ where researchers are able to upload their study results for community sharing, we recommend that researchers upload their effect estimate and uncertainty data, in addition to (or instead of) just statistical values.

## 5 Conclusions

Three aspects of information waste are involved in the conventional neuroimaging data analysis: a) the implicit adoption of the principle of insufficient reason in massively univariate analysis, b) hard dichotomization through multiple testing adjustment, and c) sole focus on statistical evidence without revealing effect magnitude and the associated uncertainty. Under the Bayesian multilevel framework, the data hierarchy across space can be captured and regularized to prevent overfitting and information waste. In addition, full results are available without artificial dichotomization. For future analyses, one may consider the following three aspects regardless of the modeling framework: 1) avoid hard thresholding; 2) report results that contain both estimate magnitude and uncertainty; 3) incorporate data hierarchies into modeling.

## 6 Acknowledgments

GC, PAT, and RWC were supported by the NIMH and NINDS Intramural Research Programs (ZICMH002888) of the NIH/HHS, USA. JS was supported by the National Institute of Mental Health (K23MH113731). LP was supported by the National Institute of Mental Health (MH071589 and MH112517).

## Footnotes

a See https://afni.nimh.nih.gov/pub/dist/doc/htmldoc/tutorials/meta/basic_bml.html for the example data and short `R` code used to perform this example meta-analysis.

b For example, a zero variance estimate (*τ*^2^ = 0) may arise under the conventional framework, especially when the number of studies is small. Such an implausible boundary estimate would not occur under the Bayesian formulation^23^.

